# The fasciola cinereum subregion of the hippocampus is important for the acquisition of visual contextual memory

**DOI:** 10.1101/748301

**Authors:** Seong-Beom Park, Eun-Young Lee, Heung-Yeol Lim, Seung-Woo Yoo, Eunsoo Lee, Hyun-Suk Jung, Woong Sun, Inah Lee

## Abstract

The fasciola cinereum (FC) is a subregion of the hippocampus that has received relatively little research attention compared with other hippocampal subregions with respect to anatomical characteristics and functional significance. Here, we show that the FC exhibits clear anatomical borders with CA1. The FC consists of granule cells, but adult neurogenesis was not found. The FC receives inputs from the lateral entorhinal cortex and perirhinal cortex while projecting to the crest of the dentate gyrus (DG). Neurotoxic lesioning of the FC using colchicine impaired the acquisition, but not retrieval, of visual contextual memory in rats. FC lesions also impaired place recognition and object-in-place memory. Place cells in the FC showed robust place fields but fired only transiently in their fields compared with those in CA1. Our findings suggest that the FC may play critical roles in learning a novel environment by facilitating pattern separation in the DG of the hippocampus.

## Introduction

The hippocampus is important for spatial navigation(Morris et al., 1982) and episodic memory(Scoville and Milner, 1957). Computational models have been put forth to explain the functions of its major subregions, including the dentate gyrus (DG), CA3, CA2, and CA1(Marr, 1971, Treves and Rolls, 1992, Lee et al., 2004). However, one hippocampal subregion, the fasciola cinereum (FC), has been almost ignored in the literature, especially with respect to its functional contributions to hippocampal information processing.

Only a handful of anatomical studies have been conducted on the FC(Arnold, 1838, Hill, 1895),(Swanson, 2014). Early studies reported that the FC is composed of granule cells(Hjorth-Simonsen and Laurberg, 1977) or pyramidal cells(Ganser, 1882, Hjorth-Simonsen and Zimmer, 1975). It has also been reported that the FC receives inputs from the lateral entorhinal cortex (LEC) in rats(Hjorth-Simonsen, 1972). Some similarities in genetic profiles between the FC and CA2 in mice have also been reported (**Table S1**). However, there is confusion in the literature as to the location of anatomical boundaries between the FC and CA1. In rodents, some anatomical atlases delineate the boundaries of FC in the form of a band-shaped structure located at the medial tip of the hippocampus near the midline of the brain(Swanson, 2018, Paxinos and Watson, 2009), whereas other studies extend its lateral borders to include the distal-most part of CA1(Boccara et al., 2015, Henriksen et al., 2010). Furthermore, whether the FC is an independent subregion or an extension of another hippocampal subregion, such as the DG(Hjorth-Simonsen, 1972) or CA2(Laeremans et al., 2013), has been a matter of controversy.

In the current study, we characterized the FC anatomically to address the abovementioned issues. We also investigated the physiological characteristics of principal neurons in the FC and their possible functions in hippocampal learning and memory, which, to our knowledge, has never been reported in the literature. Our findings show that the anatomical border between FC and CA1 can be clearly defined by various criteria, including genetic profile, input-output connectivity, and cellular morphology. The FC projects only to the DG in the hippocampus, and rats with selective lesions in the FC were impaired with respect to acquisition of novel visual contextual memory compared with controls, but not in retrieving familiar visual contextual memories. Place cells were found to be present in the FC, but they exhibited some important differences compared with place cells in CA1. Converging evidence from anatomical, behavioral, and physiological findings of our study strongly suggest that the FC is an anatomically discrete and functionally critical subregion of the hippocampus.

## Results

### The FC as a distinct subregion of the hippocampus with clear boundaries

To investigate whether the FC is a part of other hippocampal subregions or is an independent region, we first defined the boundaries between the FC and CA1 (**Fig. 1a**). Consistent with prior reports(Haug, 1974), the FC subregion was clearly delineated by its sharp borders in Timm-stained sections (**Fig. 1b**). The abrupt changes in cell density in Nissl-stained sections also clearly showed the FC boundaries (**Fig. 1b**). The border between the FC and CA1 was also marked by immunostaining for RGS14 (regulator of G-protein signaling 14), a genetic marker for both the FC and CA2Lein et al. (2005), and WFS1 (Wolfram syndrome 1), a marker for CA1(Takeda et al., 2001) (**Fig. 1c**; **Table S2**). RGS14 was expressed in both the FC and CA2, whereas WFS1 immunostaining was detected in both the distal and distal-most subdivisions of CA1 (dCA1 and dmCA1, respectively), matching the borders identified in Nissl- and Timm-stained sections.

**Figure 1.**
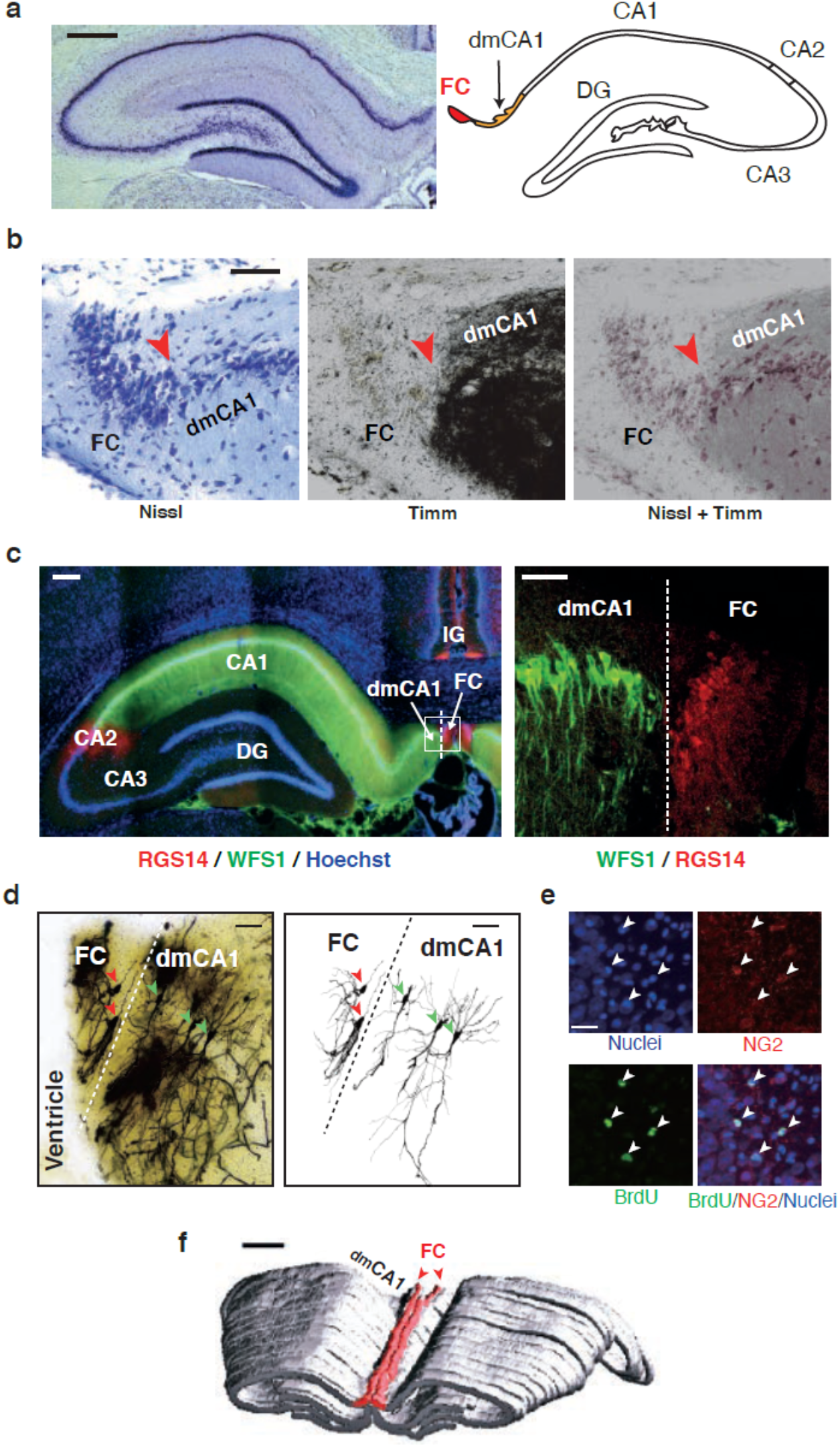
Anatomical characteristics of the FC. **a**. The FC subregion of the hippocampus. The FC and the distalmost CA1 (dmCA1) are shown as Nissl-stained tissue (*left*) and a schematic diagram (*right*). Scale bar: 500 *μ*m. The boundary between the dmCA1 and dCA1 is marked based on a previous study(Boccara et al., 2015). **b**. Boundary between the FC and dmCA1. The lateral border of the FC is clearly visible (red arrowhead) in thionin-stained (*left*), Timm-stained (*middle*), and merged (*right*) sections. Scale bar: 50 *μ*m. **c**. Differential gene expression across hippocampal subregions. *Left:* Expression of RGS14 (red) was only detected in the FC and CA2, whereas WFS1 expression (green) was only found in CA1, including the dmCA1; nuclei are stained with DAPI (blue). Scale bar: 250 *μ*m. *Right:* Magnified view of the white-boxed area. Scale bar: 50 *μ*m. **d**. Morphological differences in neurons between the FC and CA1. FC cells (red arrowheads) and dmCA1 cells (green arrowheads) are visualized in a photomicrograph of Golgi-Cox–stained tissue (*left*) and in individually traced and reconstructed images (*right*). Scale bar: 50 *μ*m. **e**. BrdU-labeled cells in the FC are co-localized with NG2-labeled glial cells (oligodendroblasts; arrowheads). Scale bars: 25 *μ*m. **f**. Three-dimensional view of the FC in the hippocampus. The FC subregion is highlighted in red (arrowheads). Scale bar: 1 mm.

Because the FC and CA2 share some genetic markers, the FC was once considered an extension of CA2(Laeremans et al., 2013). However, unlike CA2, cells in the FC stained using the Golgi-Cox method (**Fig. 1d**, red arrowhead; **Fig. S1**) are characterized by round cell bodies from which dendritic branches fan out (extending only toward the ventricular surface)—common features of granule cells in the DG(Andersen, 2007). In contrast, cells in the dmCA1 (**Fig. 1d**, green arrowheads) showed morphological characteristics typical of pyramidal neurons in CA1, such as noticeable apical and basal dendrites and pyramidal cell bodies.

To examine the possibility that the FC might be an extension of the DG(Hjorth-Simonsen, 1972), we tested whether adult neurogenesis occurred in the FC as it does in the DG(Altman and Chorover, 1963). For this purpose, we injected BrdU, a marker of neurogenesis, and allowed 7 d for labeling of adult-born cells; we then assessed expression after 3 wk. We found BrdU-labeled neurons (BrdU^+^/NeuN^+^ cells) in the DG but not in the FC (**Fig. S2a** and **2b**). Although BrdU^+^ cells were observed sparsely in the FC, further analyses revealed that these cells are glial cells and not neurons, as evidenced by both expression of the oligodendroblast marker NG2^+^ and the absence of NeuN expression (**Fig. 1e**; **Fig. S2b** and **2c**). These results suggest that the FC is distinct from the DG and should be treated as an independent subregion of the hippocampus.

The border between these different cell types matched those identified by Nissl-staining, Timm-staining, and immunostaining methods. These results unmistakably identify distinct boundaries between the FC and dmCA1. A three-dimensionally reconstructed image of the dorsal hippocampus showed the FC running longitudinally along the entire dorsal hippocampus (**Fig. 1f**).

### Afferent and efferent connectivity of the FC

Since the anatomical connectivity of the FC has rarely been studied, we injected Retrobeads (RBs), a retrograde tracer, into the FC (**Fig. 2a**). RB retrograde labeling was detected mostly in the parahippocampal region, including layer II of the LEC and layer II/III of the perirhinal cortex (PER) (**Fig. 2a** and **2b**). In the posterior LEC, RBs were found in deeper layers adjoining the borders with the postrhinal cortex (POR) (**Fig. 2a**; **Fig. S3a**). Sparse RB labeling was detected in the lateral supramammillary area (**Fig. 2a**; **Fig. S3b**). Interestingly, an examination of the FC along the longitudinal axis revealed the presence of RBs in other areas of the FC into which RBs were not directly injected (**Fig. 2b**), suggesting that cells in the FC are intrinsically interconnected with each other, unlike in CA1.

**Figure 2.**
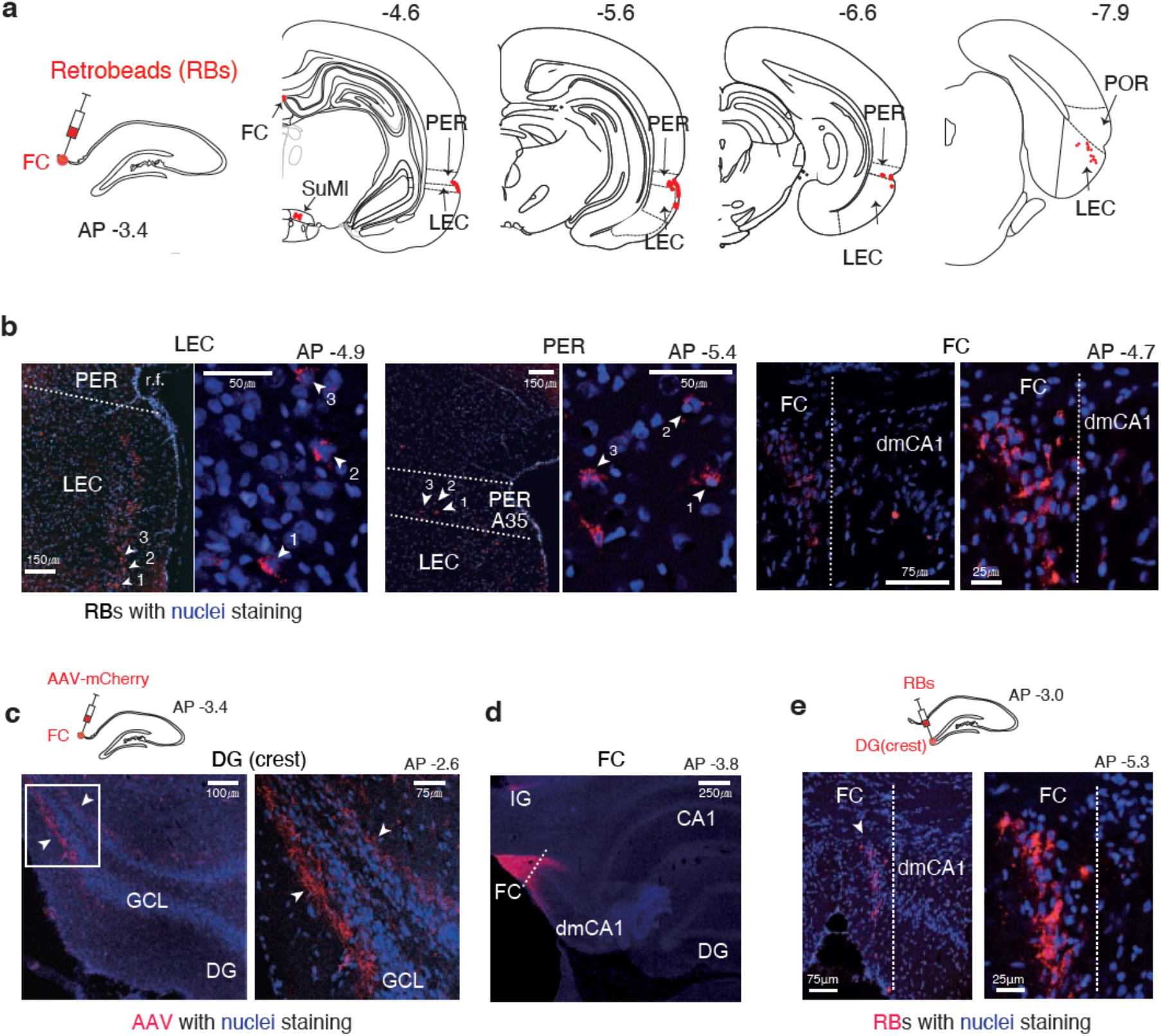
Anatomical connectivity of the FC. **a.** Afferent regions of the FC. Retrograde tracing for the afferent regions of the FC was conducted by injecting Retrobeads (RBs) into the FC. Locations with RB-containing cells (red dots) were marked on the atlas (modified from the Paxinos and Watson; parahippocampal boundaries are marked according to Burwell(Paxinos and Watson, 2009, Burwell, 2001)). Numbers indicate the relative positions of sections posterior to bregma (in mm). RBs were detected mostly in layer II of the LEC and layer II-III in the PER (A35). Posteriorly, RBs were found in deeper layers of the LEC adjoining the borders with the POR. SuMl, lateral supramammillary area. **b**. Retrograde labeling of cells projecting to the FC. Nuclei (Hoechst staining) and RBs are shown in blue and red, respectively. Cells projecting to the FC were found in the LEC (*left*) and PER (*middle*). Intrinsic connections within the FC were also found (*right*). dmCA1, distalmost CA1; rf, rhinal fissure. **c**. FC projections to the DG. Axons of FC cells containing AAV-mCherry (red; injected into the FC) were detected in the molecular layer (ML) of the crest of the septal DG. White arrowheads indicate axons immediately adjacent to the granule cell layer (blue; stained with DAPI). GCL, granule cell layer. **d.** Anterograde transport of AAV-mCherry within the FC itself via intrinsic connections was also detected. IG, indusium griseum. **e.** RBs injected into the crest of the DG were transported retrogradely to the FC.

To identify the efferent targets of the FC, we injected an adeno-associated virus expressing the fluorescent marker mCherry (AAV-CamKIIa-mCherry) or enhanced green fluorescent protein (AAV-CamKIIa-EGFP) into the FC. Excitatory axons of the FC were found in the molecular layer of the crest region of the DG near the septal tip (**Fig. 2c**) and within the FC itself (**Fig. 2d**). The FC also projected to the septohippocampal nucleus, septofimbrial nucleus, indusium griseum and the septal tip of the Cornu Ammonis, but did not project to CA1 (**Fig. S3c–3e**). RBs injected into the crest of the DG were found in the FC along the longitudinal axis, confirming the results of our anterograde tracing study (**Fig. 2e**).

The borders between the FC and CA1 were also confirmed by injecting RBs into the DG and subiculum, the major output areas of the FC and CA1, respectively (**Fig. 2e**; **Fig. S3f**). Specifically, RBs injected into the crest of the DG were found in the FC, whereas RBs injected into the subiculum were only found in CA1 (up to dmCA1) and not in the FC. Taken together, these observations suggest that, among currently known hippocampal subregions, the FC is the only subregion in the hippocampal formation that sends its major outputs exclusively to the DG.

### The FC is important for the acquisition, but not retrieval, of visual contextual memory

Given the prominent efferent connections from the FC to the DG identified in our study, we tested the functional roles of the FC in a visual contextual memory (VCM) task, which requires intact DG function in rats(Ahn and Lee, 2014, Lee and Lee, 2020). For this purpose, we developed an FC-lesion model using colchicine, which is known to fairly selectively eliminate the DG, leaving other subregions largely unaffected(Walsh et al., 1986). We successfully ablated the FC using colchicine (**Fig. 3a**; **Fig. S4** and **5**; **Table S3**), presumably because of the similar cell type (i.e., granule cells) found in both the DG and FC.

**Figure 3.**
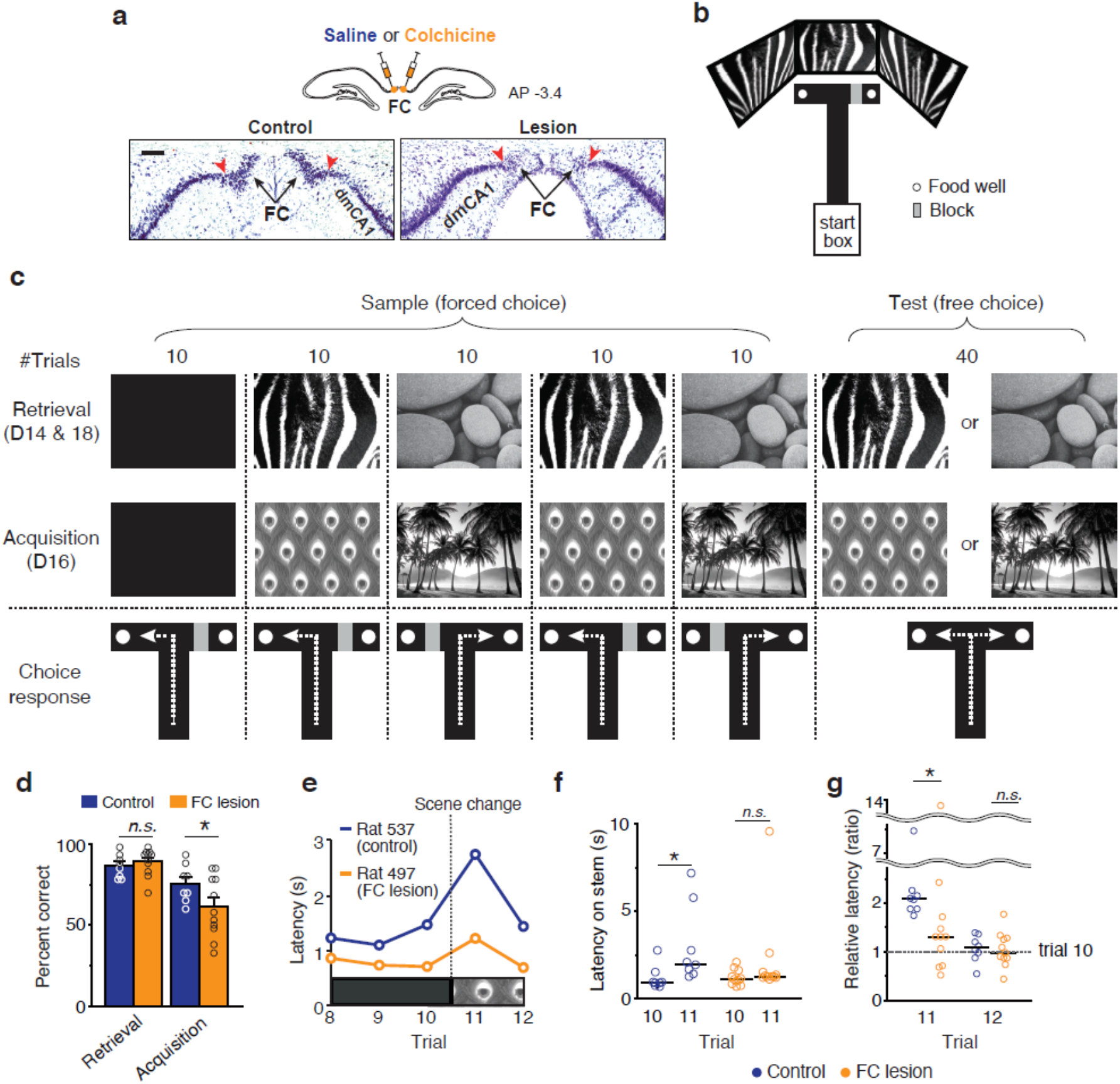
Lesions of the FC impair acquisition, but not retrieval, of contextual memory. **a.** Selective lesioning of the FC with colchicine. Injection sites (*upper*) and histological confirmation of lesions of the FC (*lower*; Nissl staining) are shown. Red arrowheads mark the boundaries between the FC and dmCA1, and black arrows indicate the FC region. Scale bar: 100 *μ*m. **b.** Behavioral apparatus. The gray rectangle represents a transparent acrylic block used to restrict access to the arm. **c.** Experimental schedule and behavioral paradigm. Rats were trained with familiar scenes before surgery. After a recovery period (14 d), rats were tested with the same scenes for 1 d (D14, *retrieval*) and then experienced two more sessions with new scenes (D16, *acquisition*). They were tested again with the originally trained old scenes (D18, *retrieval*). One day of rest was given between testing. In a given session, the rat experienced 90 trials in total (50 sample trials and 40 test trials). Sample trials were grouped into five blocks (10 trials per block). **d.** Deficits in the acquisition, but not retrieval, of scene memory in the FC-lesion group. **e-g.** Differential response to the first introduction of the novel scene (peacock feather pattern in this example) between control and lesion groups. A representative example showing the difference in latency between two rats, one from the control group and the other from the FC-lesion group, in the trial in which the novel visual scene was first presented (**e**). The average latency (**f**) and relative latency compared to trial 10 (**g**) for all rats are also shown. Statistical results are described in Table S4. Data represent means ± SEM for parametric tests (**d**) and median for non-parametric tests (**f-g**). Individual data points are also shown. (*P < 0.05).

In a modified version of the VCM task, rats were required to make a forced turn (left or right) on a T-maze while a patterned visual stimulus (i.e., visual context or scene) associated with each response was displayed in LCD monitors around the choice arms of the maze (**Fig. 3b** and **3c**). Rats experienced the two visual contexts twice in an alternate blocked fashion (10 trials per block, two blocks for each visual context; *sample* blocks). Then, during a *test* block (40 trials), in which one of the visual contexts was displayed, the rat was allowed to freely choose one of the arms (**Fig. 3c**). Rats were trained to the criterion of >75% correct for each visual context in the testing session, after which a colchicine lesion or sham lesion (control) was made in the FC (control, n = 8; lesion group, n = 11; **Table S3**). After allowing 2 wk to recovery from surgery, rats were tested every other day, beginning on day 14 (D14), with pre-surgically trained familiar visual contexts (*retrieval*) followed by novel contexts (*acquisition*) on D16. Retrieval of familiar contextual memory was also tested again on D18 (**Fig. 3c**).

Rats retested with the originally trained, thus familiar, stimulus after recovery from surgery (D14) showed no significant difference in performance between control and lesion groups (P = 0.654, two-way mixed ANOVA with Bonferroni correction) (**Fig. 3d**; **Table S4**). However, the performance of the lesion group was significantly impaired compared with controls during the acquisition of novel contextual behavior with presentation of new visual contexts (P = 0.019, two-way mixed ANOVA with Bonferroni correction) (**Fig. 3c**; **Table S4**). No significant difference was found in latency between the two groups (t_(17)_ = 0.5, P = 0.592, t-test) (**Table S4**). The normal performance of the FC-lesion group during memory retrieval on D14 was also verified in retests of previously learned contextual behavior (retrieval) on D18 (t_(17)_ = 0.5, P = 0.625, t-test) (**Table S4**).

In our VCM task, rats tended to pause on the stem of the T-maze when a new visual stimulus was first encountered; this usually made the latency of that particular trial longer than that in other trials in which stimuli became more familiar. Interestingly, there was a significant difference in latency between the control and lesion groups (Z = 2.4, P = 0.017; Mann-Whitney U test) on the first trial (trial 11) in which a novel visual stimulus was presented for the first time (between trials 10 and 11: Z = 2.5, P = 0.012 for controls; Z = 0.9, P = 0.374 for the FC-lesion group; Wilcoxon Signed rank test) (**Fig. 3e** and **3f**; **Table S4**). This phenomenon occurred only when the rat saw the novel context for the first time on trial 11 (**Fig. 3f**), and not on subsequent trials—not even the next trial (**Fig. 3g**). That is, no significant difference in latency was observed on trial 12 between control and lesion groups (Z = 0.3, P = 0.741, Mann-Whitney U test) (**Fig. 3g**).

Our findings suggest that the FC plays key roles in learning a new visual context and its associated behavior during the very early phase of acquisition. Furthermore, based on our series of object-manipulated experiments, the FC was found to be essential in recognizing objects with their associated places (or background contexts; **Fig. S6a-6c**), but not in simple object recognition (**Fig. S6d** and **S6e**).

### Physiological properties of single units in the FC

To examine whether principal cells in the FC show similar physiological characteristics compared with neurons in other subregions of the hippocampus, we recorded single units from the FC while the rat foraged for randomly scattered food in a square environment. To avoid damaging the superior sagittal sinus, we implanted a bundle carrying twenty-four tetrodes angled 20° medially and a hyperdrive angled 10° laterally from the vertical axis (**Fig. 4a**). Histological results showed that the tetrodes targeted the FC, dmCA1, and dCA1 (**Fig. 4a**). Compared to cell recordings in CA1, we found it very difficult to record single units from the FC because the structure is very tiny and narrow (approximately 80 × 230 *μ*m in a coronal section), and the recording tetrodes must approach the FC obliquely to avoid damaging the superior sagittal sinus. Nevertheless, histological results showed that multiple tetrodes accurately targeted the FC (n = 14) as well as the dmCA1 (n = 45) and dCA1 (n = 23) (**Fig. 4a**). Ultimately, putative complex spiking neurons were recorded from the FC (n = 121), dmCA1 (n = 369), and dCA1 (n = 240).

**Figure 4.**
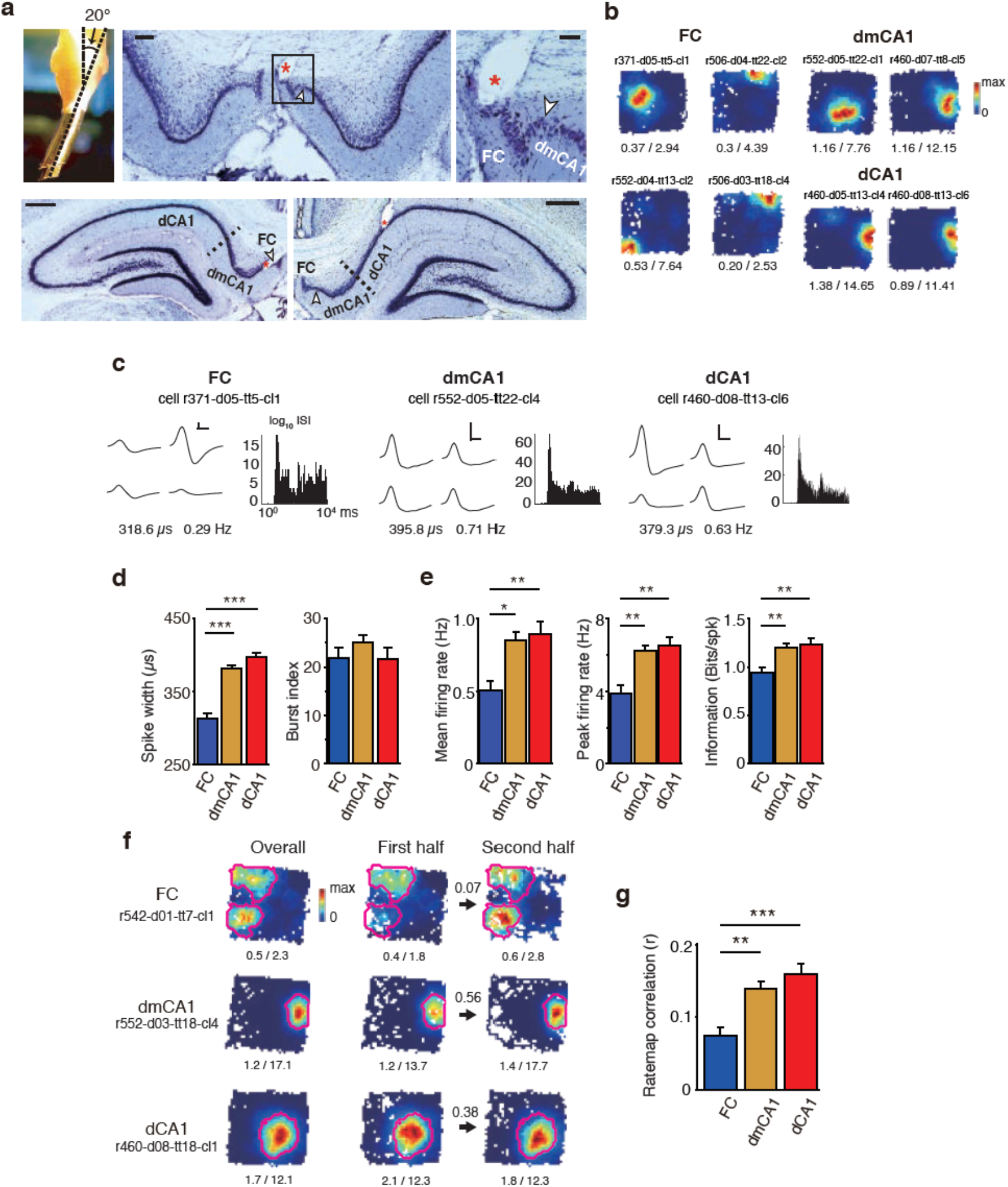
Physiological properties of cells in the FC and CA1. **a.** Recording electrode bundle and histological verification of tetrode tips. An angled bundle (20° medially) carrying 24 tetrodes was installed in the hyperdrive (*upper left*). Representative example of an FC tetrode track (*upper middle*). Magnified view of the tetrode tip area (*; *upper right*). Tetrode tracks showing the final recording positions in dmCA1 (*lower left*) and dCA1 (*lower right*) regions are also shown. Scale bars: 100 *μ*m (*upper middle*), 50 *μ*m (*upper right*), and 500 *μ*m (lower panels). **b.** Representative place fields of principal cells in the FC, dmCA1, and dCA1 in a square arena. The numbers below are mean (*left*) and peak (*right*) firing rates (Hz) of the cells. **c.** Waveforms of representative cells. Representative waveforms (*left*) and inter-spike interval (ISI) histograms (log scale, *right*) of single units recorded from the FC, dmCA1, and dCA1. Numbers below the plots indicate the spike width (*left*) and firing rate (*right*) during rest. Vertical scale bar, 100 *μ*V; horizontal scale bar, 200 *μ*s. **d**. Comparison of non-spatial firing characteristics (spike width and burst index) among the FC, dmCA1, and dCA1 (means ± SEM). **e.** Comparison of the mean firing rate, peak firing rate, and spatial information among the FC, dmCA1, and dCA1 during foraging. **f**. Representative examples of place fields recorded from the FC, dmCA1, and dCA1. Rate maps for the entire session (15 min) are shown on the left (*overall*). Rate maps for the first half of the session (*first half*) and the second half of the session (*second half*) are shown on the right. The numbers below the rate maps are mean and peak firing rates (Hz). The stability of the rate map (correlation coefficient) between the first and second halves of the session is denoted above the arrow. Magenta polygons indicate the boundaries of place fields. **g**. Comparison of rate-map stability, measured by spatial correlation (r), between the first half and second half of the session. Statistical results are described in Table S6. Data represent means ± SEM (*P < 0.05, **P < 0.01, ***P < 0.001).

During foraging, cells in the FC fired in a place-specific manner. Forty cells in the FC (33%), 170 cells in dmCA1 (46%), and 104 cells in dCA1 (43%) were classified as place cells. Rate maps of place cells recorded from the FC and CA1 were almost indistinguishable by visual inspection (**Fig. 4b**). However, we found some notable differences in firing properties of neurons between the two subregions. Specifically, spikes recorded from cells in the FC exhibited narrower waveforms than those in CA1 (F_(2, 311)_ = 40.8, P < 0.001; FC vs. dmCA1, P < 0.001; FC vs. dCA1, P < 0.001; dmCA1 vs. dCA1, P = 0.017; one-way ANOVA with Bonferroni correction) (**Fig. 4c** and **4d**; **Table S6**). Cells in both the FC and CA1 showed similar bursting spiking patterns (F_(2,311)_ = 1.0, P = 0.37, one-way ANOVA) (**Fig. 4d**; **Table S6**). Moreover, the mean firing rate was significantly lower in the FC than in CA1 (F_(2,311)_ = 3.7, P = 0.025; FC vs. dmCA1, P = 0.014; FC vs. dCA1, P = 0.009; dmCA1 vs. dCA1, P = 0.643; one-way ANOVA with Bonferroni correction), as was the peak firing rate (F_(2,311)_ = 5.4, P = 0.005; FC vs. dmCA1, P = 0.003; FC vs. dCA1, P = 0.002; dmCA1 vs. dCA1, P = 0.598; one-way ANOVA with Bonferroni correction) (**Fig. 4e**; **Table S6**). Individual spikes of FC cells carried significantly less spatial information than those of CA1 cells (F_(2,311)_ = 4.2, P = 0.016; FC vs. dmCA1, P = 0.009; FC vs. dCA1, P = 0.006; dmCA1 vs. dCA1, P = 0.632; one-way ANOVA with Bonferroni correction) (**Fig. 4e**; **Table S6**).

Interestingly, spatial representations of FC cells were significantly less stable than those of place cells in CA1, as reflected in the significantly lower spatial correlation between rate maps for the first and second halves of the recording session in the FC compared with CA1 (F_(2,311)_ = 5.9, P = 0.003; FC vs. dmCA1, P = 0.005; FC vs. dCA1, P <0.001; dmCA1 vs. dCA1, P = 0.254; one-way ANOVA with Bonferroni correction) (**Fig. 4f** and **4g; Table S6**). Importantly, FC cells exhibited significantly different firing patterns across all criteria indicated above compared with cells in the dmCA1, which is immediately adjacent to the FC anatomically. Moreover, we found no significant difference in these measures between the dmCA1 and dCA1, thus physiologically verifying the anatomical borders delineating CA1 and the FC in our study (**Table S6**). In addition to assessing spatial firing properties of place cells, we also examined whether robust sharp-wave ripple (SWR) activity—a characteristic rhythm in most subregions of the hippocampus(Buzsaki, 2015, Swaminathan et al., 2018)—was detectable in the FC. To this end, we simultaneously recorded local field potentials (LFPs) and spiking activities of place cells in both the FC and CA1 during sleep (**Fig. 5a**). Strong SWR events were observed in both the dmCA1 and dCA1 (**Fig. 5b** and **5c; Fig. S7**). In contrast, LFP power at the ripple band (150-250 Hz) in the FC during SWR events of CA1 was significantly weaker than that in CA1; in fact, it was similar to that recorded from the corpus callosum (F_(3,292)_ = 132.5, P < 0.001; FC vs. dmCA1, P < 0.001; FC vs. dCA1, P < 0.001; dmCA1 vs. dCA1, P = 0.656; FC vs. CC, P = 0.384, one-way ANOVA with Bonferroni correction) (**Fig. 5d**; **Table S7**). Also, place cells in CA1 showed prominent bursting activity during SWR events (**Fig. 5e**); this activity was quantified by dividing the firing rate during SWR events by the firing rate during the rest of the recording period (non-SWR) (**Fig. 5f**). However, place cells in the FC showed significantly less bursting activity during SWR events (**Fig. 5e; Fig. S7**) than those in CA1 (F_(2,247)_ = 13.447, P < 0.001; FC vs. dmCA1, P < 0.001; FC vs. dCA1, P < 0.001; dmCA1 vs. dCA1, P = 0.088, one-way ANOVA with Bonferroni correction) (**Fig. 5f**; **Table S7**). These results collectively confirm the anatomical boundaries between the FC and CA1 in the current study and further support the conclusion that the FC is an independent subregion of the hippocampus.

**Figure 5.**
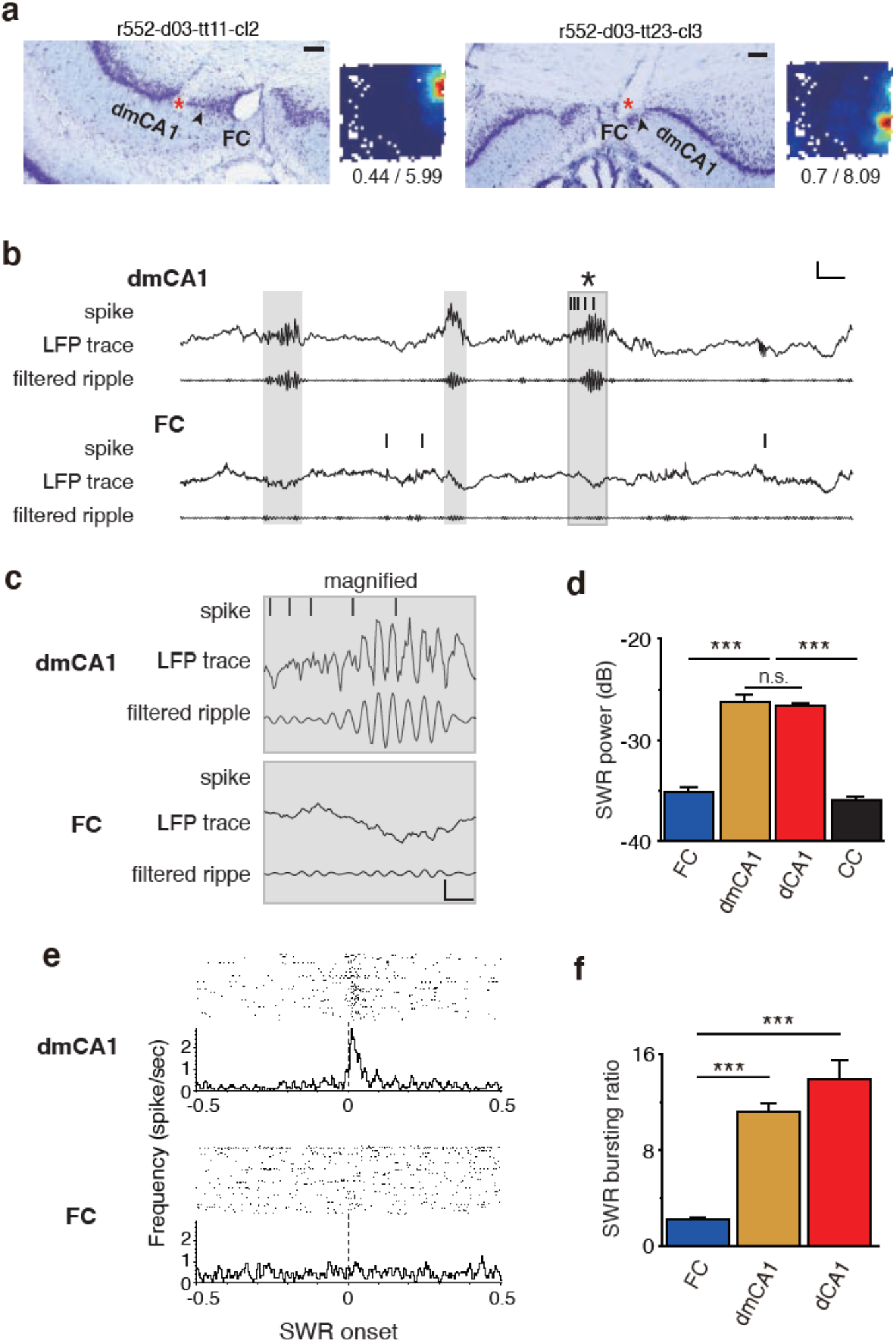
SWR-related spiking activities in CA1 and the FC. **a**. Histological verifications of the tips (red asterisk) of tetrodes and the firing rate maps of simultaneously recorded cells (from the tetrodes shown) in both the dmCA1 (*left*) and FC (*right*). Numbers below the firing rate map indicate mean and peak firing rates (Hz). Black arrowhead denotes the border between the dmCA1 and FC. Scale bar: 100 *μ*m. **b**. Spike train and LFP trace for both units shown in (**a**) during the post-sleep session after the foraging session. Raw and band-pass filtered LFPs (150-250 Hz) are shown. The gray zone denotes the SWR event period. Vertical scale bar, 400 *μ*V; horizontal scale bar, 50 ms. **c**. Magnified view of the last SWR event in **(b)**. Vertical scale bar, 200 *μ*v; horizontal scale bar, 10 ms. **d**. Power of ripple oscillation detected in the tetrodes in (**a**). **e**. Peri-event time histogram (PETH, *lower*) and raster plot (*upper*) aligned to SWR onset timing for the cells presented in (**b**). **f**. SWR bursting ratio, defined as the average firing rate within the SWR period divided by the mean firing rate of ripple events in the subregion. Statistical results are described in Table S7. Data represent means ± SEM (*P < 0.05, **P < 0.01, ***P < 0.001).

### Stronger rate remapping for visual contextual change by place cells in the FC than in CA1

How could the FC contribute to hippocampal learning and memory? Considering that the FC sends its major output only to the DG in the hippocampus, we recorded single units from the FC while the rat performed the VCM task(Ahn and Lee, 2014, Lee and Lee, 2020)—the same task previously used for examining the effects of FC lesions (**Fig. 3**). We have demonstrated that, in the VCM task, the DG is necessary for CA3 cells to exhibit pattern-separating signals between different visual contextual environments(Lee and Lee, 2020). In the modified version of the task, rats were trained with familiar visual contexts before receiving hyperdrive-implanting surgery. Once tetrodes reached the target regions, rats were tested with familiar visual contexts (zebra stripes and pebbles) on days 1 and 3, and with novel visual contexts on day 2 (peacock and palm trees) and day 4 (bamboos and mountains).

Performance of rats (n = 8) significantly exceeded chance level during retrieval (D1 and D3; 91.3 ± 1.8; t_(7)_ = 22.9, P<0.001, t-test) and acquisition (D2 and D4; 84.5 ± 2.9, mean ± SEM; t_(7)_ = 11.8, P < 0.001, t-test), and there was no significant difference in performance between the two learning stages (t_(14)_ = 2.0, P = 0.070, t-test) (**Fig. 6a**, right). Recordings of place cells from the FC (n = 32) and dmCA1 (n = 148) while rats (n = 4) performed the VCM task showed that place cells in the dmCA1 exhibited rate remapping for different visual contexts, as reported previously(Delcasso et al., 2014, Lee and Lee, 2020, Lee et al., 2018) (**Fig. 6b**), and revealed that cells in the FC exhibited similar behavior (**Fig. 6c**). Measurement of the amount of rate remapping using a *rate modulation index (RMI)* showed that cells in the FC exhibited higher RMI scores than those in the dmCA1, and neither the learning stage (acquisition vs. retrieval) nor its interaction with the subregion significantly explained the difference in RMI scores between the FC and dmCA1 (subregion, F_(1,128.4)_ = 4.1, P = 0.044; learning stage, F_(1,128.4)_ = 0.3, P = 0.589; subregion × learning stage, F_(1, 128.4)_ = 0.0, P = 0.912; two-way ANOVA) (**Fig. 6d**; **Table S8**). We also verified that the differences in basic firing properties of place cells between the FC and dmCA1 previously observed during random foraging (**Fig. 4**) were similarly observed in the VCM task (all P-values < 0.05; **Table S8**). Again, no significant difference was found between the two subregions with respect to learning stage or its interaction with the subregion (**Table S8**).

**Figure 6.**
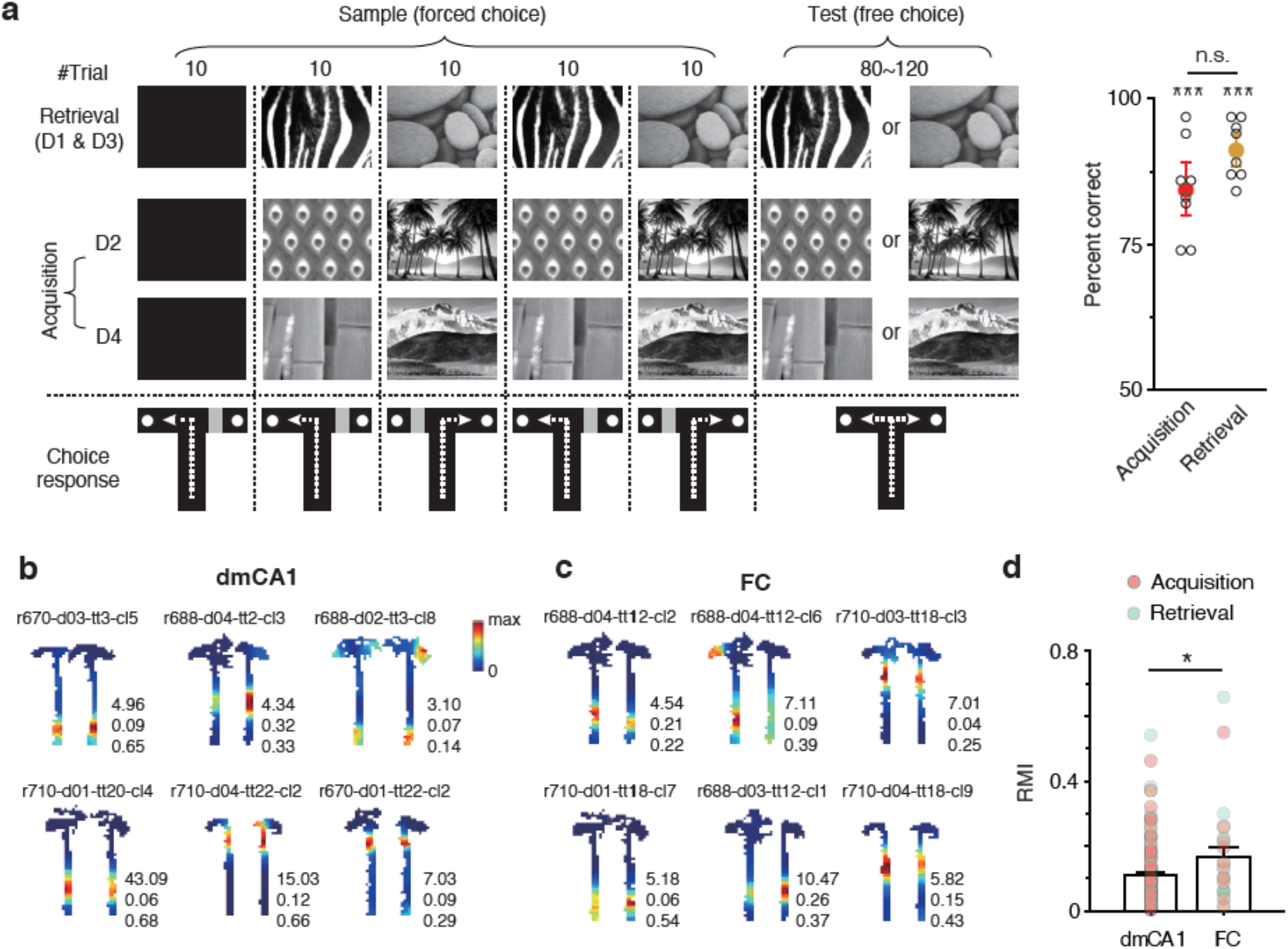
Visual contextual modulation of firing in FC place cells in the VCM task. **a**. Experimental schedule and behavioral paradigm (*left*). The task was similar to that used for the lesion study (Figure 4) except for the presence of an additional acquisition session (D4) and an increased number of trials for physiological recordings during testing. Overall performance during acquisition (D2 and D4) and retrieval (D1 and D3) (*right*). **b, c**. Rate maps of place cells in the dmCA1 (**b**) and FC (**c**) for paired scene stimuli. Numbers beside rate maps indicate the peak firing rate (*top*), the amount of visual scene-dependent rate modulation (in-field rate modulation index or RMI; *middle*), and the correlation coefficient between the rate maps (*bottom*). **d**. RMI values for individual place cells in the dmCA1 and FC are shown with mean RMIs (± SEM). The colors of dots indicate that place fields were recorded during the acquisition (red) or retrieval (green) task. Place cells in the FC exhibited significantly higher rate modulation between visual contexts than those in the dmCA1. Statistical results are described in Table S8 (*P < 0.05).

### Location-specific neuronal firing is more episodic and transient in the FC compared with CA1

During random foraging, rate maps of place cells in the FC were less stable than those in CA1, based on spatial correlation measures (**Fig. 4f** and **4g; Table S6**). An examination of place fields across individual trials within a session showed that place fields of the dmCA1 remained stable after the fields had formed (**Fig. 7a**). However, place cells in the FC showed episodic firing patterns and exhibited location-specific firing only in a subset of trials within a session, as if coding place information for a limited period of time (**Fig. 7b**). That is, place fields in the FC typically appeared abruptly and remained stable only for a certain period of time. More specifically, during the VCM task, most place fields in the dmCA1 (70%) emerged in the first two recording blocks (trial 1 to 20), whereas more than half (55%) of place fields in the FC appeared after the second block (trial 21) (subregion: F_(1,178.2)_ = 6.5, P = 0.011, two-way ANOVA) (**Fig. 7c**; **Table S9**). Furthermore, most place fields in the dmCA1 (77%) remained active until the last trial, whereas approximately half (53%) of place fields in the FC disappeared before the last trial (χ^2^(1) = 12.3, P < 0.001, Chi-square test) (**Fig. 7d**). As a result, the duration of place-specific spiking activity differed between the two subregions (F_(1,180)_ = 23.0, P < 0.001, two-way ANOVA) (**Fig. 7e**; **Table S9**). An assessment of recording stability based on a comparison of differences in firing rate between pre-sleep and post-sleep stages divided by the sum of firing rates revealed no significant difference between the dmCA1 (**Fig. S8a**) and FC (**Fig. S8b**), suggesting that recordings in the FC were as stable as those in the dmCA1 (**Fig. S8c**; **Table S9**). The transient nature of location-specific firing in the FC was also verified at the population level (**Fig. S9**).

**Figure 7.**
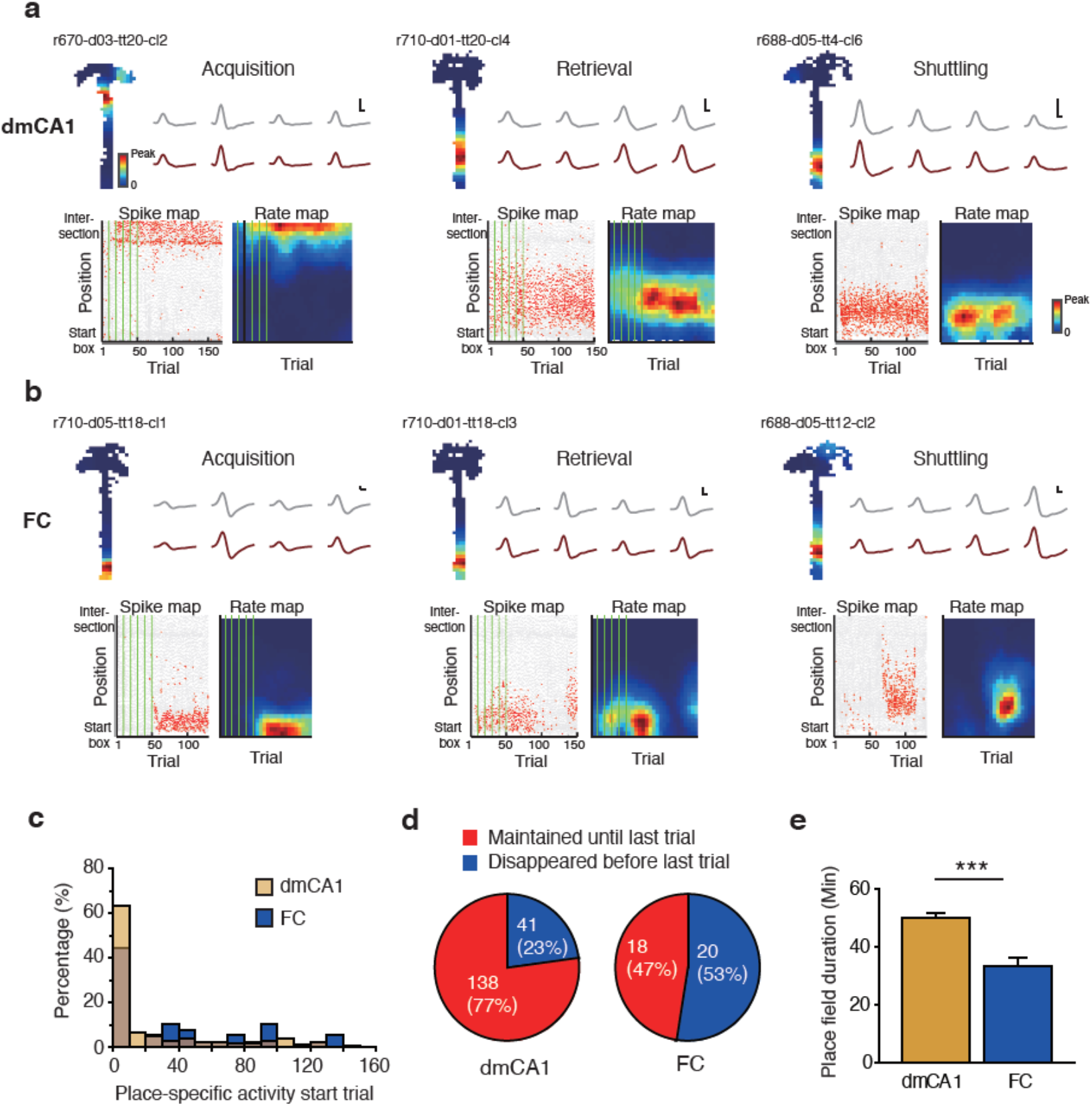
Transient spatial firing patterns of place cells in the FC. **a, b.** Place-specific firing patterns across trials and at different stages of learning in the FC (**a**) and dmCA1 (**b**). Representative rate map of a place cell recording from the FC is shown (*upper left*), with the cell’s averaged spiking waveforms recorded during pre-sleep (gray) and post-sleep (brown) stages on the right side. Horizontal scale bar, 100 *μ*s; vertical scale bar, 100 *μ*V. The same place cell’s location-specific (ordinate) spiking activities and rate map are plotted across trials (abscissa; *bottom*). Gray dots denote the position traces, and red dots indicate individual spikes of the place cell. Sample blocks are separated from each other by green vertical lines. Note that the spiking activities of FC cells appeared in a limited period compared with the sustained activities observed throughout the session in the dmCA1. **c.** Distribution of trials for which spiking activity started for individual place cells in the FC and dmCA1. Note that a higher proportion of place cells started firing at later trials in the FC compared with the dmCA1 (subregion: F_(1,178.2)_ = 6.5, P = 0.011, two-way ANOVA). **d**. Proportions of place cells that exhibited place-specific firing throughout the session (until the last trial; red) and those that quit their firing before the session was terminated (blue) in the FC and dmCA1. Note that a larger proportion of place cells stopped firing before the session ended (χ^2^(1) = 12.3, P < 0.001, Chi-square test). **e**. Mean duration of location-specific spiking activities of place cells in the dmCA1 and FC. Note the significantly smaller duration in the FC than in the dmCA1 (F_(1,180)_ = 23.0, P < 0.001, two-way ANOVA). Statistical results are described in Table S9. Data represent means ± SEM (***P < 0.001).

## Discussion

In the current study, we demonstrated that the FC subregion of the hippocampus is an independent subregion with distinct anatomical and physiological properties that differentiate it from other hippocampal subregions. The anatomical boundaries of the FC can be clearly defined using various anatomical assays. The FC exhibits unique afferent connections with distinct subregions (e.g., PER and LEC) of the parahippocampal region, and it sends its output only to the DG in the hippocampus. The FC plays an important role in encoding, but not retrieving, new visual context, as evidenced by the fact that rats lesioned in the FC showed impairment in the acquisition of contextual behavior using novel visual contexts. Furthermore, the FC plays an important role in recognizing an object in relation to its place (or context), but not in recognizing an object irrespective of its associated location. Place cells were identified in the FC and shown to modify their firing rates in response to visual contexts to a greater extent than CA1 place cells, although their spatial firing patterns were more transient and episodic than those in CA1.

### The FC is an independent subregion of the hippocampus

Prior studies sought to characterize the FC anatomically, but some inconsistencies are found in the literature. For example, some researchers defined the boundaries of the FC conservatively(Haug, 1974, Swanson, 2018), whereas others extended the boundaries more laterally(Boccara et al., 2015, Henriksen et al., 2010). In addition, some studies reported that the FC consists of granule cells(Hjorth-Simonsen and Laurberg, 1977) or pyramidal cells(Ganser, 1882, Hjorth-Simonsen and Zimmer, 1975), the former of which was confirmed in our study by the Golgi-staining method. In our study, results obtained from various histological methods converged on the conclusion that the FC is a distinct subregion that can be clearly distinguished anatomically from CA1 (i.e., the dmCA1). Importantly, the FC was found to project only to the septal DG within the hippocampus. Extending the boundaries of the FC more laterally toward the proximal CA1(Boccara et al., 2015, Henriksen et al., 2010) than the findings reported here support would inevitably shift the boundaries of the distal CA1 more laterally, potentially resulting in an erroneous functional characterization of the distal CA1. Since it has been demonstrated that the proximal and distal subdivisions of CA1 differentially receive inputs from the medial entorhinal cortex (MEC) and LEC, respectively(Steward, 1976, Sun et al., 2018), delineating the boundaries between FC and CA1 is not a trivial issue.

The FC was previously characterized in some studies as an extension of one of the hippocampal subregions, such as the DG or CA2(Laeremans et al., 2013, Hjorth-Simonsen, 1972). We found that the principal cell type in the FC was very similar to granule cells in the DG and was different from the pyramidal cells in CA subregions, including CA1, CA2, and CA3. Our results indicate that the FC is as vulnerable to colchicine as the DG based on a lesion study, and further suggest that the principal cells in the FC are granule cells(Goldschmidt and Steward, 1982). However, most genetic markers of the FC do not match those of the DG (**Table S1**), although there is some partial overlap. Most importantly, our results indicate that the major intrahippocampal target of the FC is the DG and not CA3. Taken together, our findings suggest that the FC is an independent structure that does not belong to other subregions of the hippocampus.

### The FC has place cells despite the fact that its major inputs originate from non-spatial parahippocampal regions

The traditional dual-stream theory for the medial temporal lobe (MTL) posits separate information processing streams for spatial and non-spatial information(Eichenbaum et al., 2007, Hargreaves et al., 2005). The spatial stream includes the POR and MEC, while the non-spatial stream involves the PER and LEC. Our findings show that the major inputs of the FC come largely from the PER and LEC. Therefore, according to this theory, one might expect that cells in the FC would exhibit non-spatial firing correlates, considering the firing pattern of the cells recorded from the PER and LEC(Hargreaves et al., 2005, Deshmukh et al., 2012). However, place cells were found without difficulty in the FC, and their firing patterns were as spatial as those recorded from CA1.

Our findings do not fit the traditional model and suggest that modifications of the model are in order. Specifically, recent studies have raised the possibility that parahippocampal region areas may not be as sharply segregated as once thought(Hunsaker et al., 2013, Knierim et al., 2014, Park et al., 2017, Yoo and Lee, 2017, Lee et al., 2021). Furthermore, a recent study suggested that the POR also projects heavily to the LEC and that the connectivity between the POR and MEC is not as prominent as previously thought(Doan et al., 2019). Cells in the LEC exhibit spatial correlates in relation to objects in space(Wang et al., 2018), and some LEC cells seem to code for the trace of an object’s position in space using allocentric spatial cues(Tsao et al., 2013). It was also reported that place cells in the hippocampus still show place-specific firing patterns in the absence of an object in space after inactivation or lesion of the MEC(Hales et al., 2014, Miao et al., 2015). In this case, the spatial representations of place cells in the hippocampus might be attributable to LEC inputs. The results from these prior studies suggest that place cells in the FC receive their spatial information mostly from the LEC.

### The FC is important for acquisition of new contextual memory

In the current study, we found that the FC is important for hippocampal contextual learning. Rats with FC lesions were impaired in the acquisition of novel visual context compared with controls. In particular, control rats showed hesitation behavior when they first encountered a novel visual context, but FC-lesioned rats did not. In contrast, FC-lesioned rats showed normal retrieval of old contextual memory. The contextual behavior of the FC-lesioned rat is similar to that of the DG-lesioned rat, as previously reported(Ahn and Lee, 2014), and suggests that the FC is functionally connected to the DG to detect visual contextual novelty so as to learn a novel or modified environment(Hunsaker et al., 2008).

It has been suggested that variability in neural signals is indispensable for learning and decision making(Renart and Machens, 2014). Interestingly, we found that place cells in the FC were active only during a limited time window in a recording session, whereas these cells in CA1 fired throughout the session. This transient or episodic nature of the FC allowed its population network to shift its state rapidly and dynamically compared with the CA1 network. Prior studies reported similar physiological phenomena in different areas of hippocampal memory systems. For example, neural firing patterns change over time in the LEC (Tsao et al., 2018) and CA1(Mankin et al., 2012, Ziv et al., 2013, Taxidis et al., 2020). The variability in neural activity of CA1 might be driven from CA2, in which the spatial representation of the place cell continuously changes(Mankin et al., 2015). This is likely because the temporal dynamic of time cells in CA1 and the animal’s performance in a working memory task are disrupted by inhibition of the projection from CA2 to CA1(MacDonald and Tonegawa, 2021). Although elucidating a more thorough mechanistic explanation is beyond the scope of the current study, it is possible that the episodic firing patterns of the FC may also facilitate time-sensitive contextual or place information in the hippocampus via the DG.

The dynamically changing network state of the FC may be useful for providing the DG with a dynamic contextual signal that facilitates learning and novelty detection. It is well known that place cells in the hippocampus may develop multiple place maps for the same familiar environment, sometimes even in the absence of changes in sensory inputs(Sheintuch et al., 2020), but the underlying neural mechanisms are largely unknown. Because the DG is important for pattern separation(Treves and Rolls, 1992) and is the only target of FC projections in the hippocampus, it is possible that the variability in neural activity in the FC may contribute to pattern separation of inputs coming at different times to the DG. Specifically, it has been suggested that the DG may reduce input overlap using an expansion coding scheme, with various inhibitory forces helping to achieve sparse coding(Marr, 1971, Kesner and Rolls, 2015, Nakazawa, 2017) (**Fig. S10a** and **S10b**). Neurogenesis can also contribute to pattern separation(Deng et al., 2010). Specifically, because immature granule cells show a higher firing rate than mature granule cells, they can readily respond to inputs and contribute to the formation of a new memory representation. In addition, the continuous addition of newborn granule cells in the DG may allow greater separation of the representations for temporally distant inputs(Rangel et al., 2014). However, the time course of neurogenesis would appear to be too slow to explain the pattern-separation processes in the DG for temporally adjacent inputs fed to the network on a time scale of seconds to hours(Deng et al., 2010, GoodSmith et al., 2017). The episodic firing patterns of the FC in our study may facilitate pattern separation of temporally adjacent contextual inputs in the DG by providing transient inputs (**Fig. S10c**).

## Acknowledgments

This study was supported by the National Research Foundation of Korea (NRF 2017M3C7A1029661, 2018R1A4A1025616, 2019R1A2C2088799) and the BK21 FOUR by the Ministry of Education and NRF.

## Author contributions

Conceptualization - SBP, HSJ, IL; Methodology - SBP, HSJ, EL, WS, IL; Investigation: SBP, EYL, HYL, SWY, EL, IL; Visualization - SBP, EL, WS, IL; Supervision - SBP, WS, IL; Writing and editing - SBP, IL

## Data and materials availability

Most data are available in the main text or the supplementary materials, and other data not shown are available upon request.

## Materials and Methods

### Subjects

Male Long-Evans (LE; n = 70) and Sprague-Dawley (SD; n = 54) rats, individually housed with a 12-12 h light-dark cycle, were used in the study. All behavioral experiments were conducted during the light cycle. All protocols were approved by the Institutional Animal Care and Use Committee of the Seoul National University.

### Golgi staining

Three rats were sacrificed by inhalation of an overdose of CO_2_, and their brains were extracted without perfusion. Brains were separated into 5-mm blocks that included the dorsal hippocampus and were subsequently processed according to the guideline of the FD Rapid GolgiStain Kit (FD Neuro Technologies, Columbia, MD, USA). Brains were frozen and serially cut into 200-µm thick sections using a sliding microtome (HM 430; Thermo Fisher Scientific, MA, USA) and mounted on gelatin-coated microscope slides. Sections were washed with distilled water two times before staining and two times after staining (4 min per wash), stained with Solution D/E (10 min), dehydrated using an ethanol series (50%, 4 min; 75%, 4 min; 95%, 4 min; 100%, 4 × 4 min) and cleared with Histoclear II (National Diagnostics, Atlanta, GA, USA) (3 × 4 min). Slides were coverslip-mounted using Permount (Thermo Fisher Scientific), and photomicrographs were acquired using a fluorescence microscope (Eclipse 80i; Nikon) or slide scanner (MoticEasyScan One; Motic, Hong Kong).

### Anatomical tracing

SD rats (9–24 wk old) were used in the retrograde-tracing study (n = 31) and anterograde-tracing study (n = 7). For cannula implantation, rats in a stereotaxic frame were initially anesthetized with sodium pentobarbital (Nembutal, 65 mg/kg), and anesthesia was maintained using 1-2% isoflurane (Piramal, Bethlehem, PA, USA). Small burr holes were drilled, and a commercial guide cannula (26 G; Plastics One, Roanoke, VA, USA) targeting the FC, dmCA1 or DG was implanted. One or two cannulas were inserted at the following coordinates: (1) FC: −3.4 mm from bregma, 1.1-1.2 mm from the midline, and 3.7 or 4.0 mm below the dura at a 20° angle; (2) CA1: −3.4 mm from bregma, −1.2 mm from the midline, 3.0 mm below the dura; (3) DG: −3.0 mm from bregma, 2.4 mm from the midline, 4.3 mm below the dura at a 20° angle; (4) Subiculum: −5.2 mm from bregma, ±1.2 mm from the midline, 3.4 mm below the skull, and −5.8 mm from bregma, ±5.8 mm from the midline, 3.2 mm below the skull. Guide cannulas were used in tracing experiments to prevent upward spread of the tracer. Cannulas were fixed to the skull with dental cement and skull screws, after which Retrobeads (RBs) were injected through the injection cannula (33 G; Plastics One), which protruded less than 0.5 mm from the tip of the guide cannula. The injection cannula was connected to a 10 *μ*l Hamilton syringe via polyethylene tubing (PE20; Becton Dickinson, NJ, USA) and injections were controlled by a micropump (KDS-101; KD Scientific, Holliston, MA, USA). For retrograde tracing, RBs (30 nL of red or green RBs; Lumafluor, NC, USA) were injected at each injection site at a rate of 10 *μ*L/h. The rat was sacrificed 3-10 d after the injection of RBs. For anterograde tracing, AAV–CMV-mCherry (0.03 *μ*L) or eGFP(Kaspar et al., 2002) (0.05 *μ*L) (Addgene plasmid #49055; KIST, Seoul, Korea) was injected into the FC, and the rat was sacrificed 3 wk after the injection. Four rats were injected into the right FC with 0.02 *μ*L of AAV2/9-CaMKII-hChR2(E123A)-mCherry-WPRE.hGH (Catalog #AV-9-35506; Addgene plasmid #35506, Penn Vector Core, PA, USA)(Mattis et al., 2011) via a glass pipette controlled by a micropump (Legato 130; KD Scientific). The injection coordinates for the four rats were as follows: 3.5 mm posterior from bregma, 0.1 mm from the midline, and 3.6 mm below the dura (not angled) through the superior sagittal sinus.

### Histological procedures

Rats were sacrificed by inhalation of an overdose of CO_2_ and were transcardially perfused first with 0.1 M phosphate-buffered saline (PBS) and then with 4% v/v formaldehyde. The brain was extracted and fixed by soaking in a 30% sucrose-formalin at 4°C until it sank. Thereafter, the brain was coated with gelatin and fixed again in a sucrose-formaldehyde solution for 1-2 d. The brain was sectioned at 35 *μ*m for anatomical tracing and 40 *μ*m for other applications using a freezing microtome (HM 430; Thermo Fisher Scientific). Every third section was first Nissl stained (Thionin staining). Adjacent sections were used for fluorescence photomicrography and staining for nuclei (Hoechst, Cat No. 33342, Thermo Fisher, 1:1000; DAPI, H-1200, Vectashield). Photomicrography was conducted using a fluorescence microscope (Eclipse 80i, Nikon) or confocal microscope (Leica TCS SP8; Leica). For delineating region boundaries, serial sections from one LE rat were subjected to Thionin staining, Timm’s staining, and myelin staining.

### Neurogenesis

SD rats (12 wk old; n = 10) were used. BrdU was dissolved in saline solution (30-50 mg/mL) and intraperitoneally injected into rats at a dose of 100 mg/kg between 1 and 4 pm for seven consecutive days. Seven days after the last injection, five rats were transcardially perfused first with 0.1 M PBS and then with a 4% formaldehyde solution; the remaining rats were sacrificed at 21 d and perfused with 4% paraformaldehyde. Brains were post-fixed in the same fixative for 24 h and then cryoprotected in 30% sucrose, sectioned serially (40-*μ*m-thick sections), and stored in PBS containing 50% glycerol at −20°C until use. For BrdU staining, slides were immersed in 10 mM sodium citrate buffer (pH 6.0) pre-heated in a microwave for 7 min. After 5 min, the buffer (with immersed slides) was reheated for 5 min using the microwave and then allowed to cool for 30 min at room temperature. Slides were then prepared for immunohistochemistry (see below) by first washing with PBS followed by blocking. Primary antibodies against BrdU (Thermo Fisher Scientific, 1:300 or Abcam, 1:500), NeuN (Millipore, 1:500), and NG2 (Millipore, 1:500) were applied overnight at 4°C with gentle shaking. After several washes with PBS, species-appropriate fluorescence-conjugated secondary antibodies were applied for 1 h for fluorescence imaging. Sections were subsequently washed and mounted. Images were acquired using a TCS SP8 confocal laser-scanning microscope (Leica, Wetzlar, Germany).

### Immunohistochemistry

Two SD rats (8 wk old) were used for immunohistochemical marking of boundaries between the FC and CA1. Rats were anesthetized with an intraperitoneal injection of urethane and transcardially perfused with saline solution, followed by a 4% paraformaldehyde solution. The brain was extracted and stored in 4% paraformaldehyde solution until it sank to the bottom of the tube and then was transferred to a 30% sucrose-formaldehyde solution or 30% sucrose solution. Coronal sections (40 *μ*m) were frozen and cut on a sliding microtome, after which sections were incubated overnight at 4°C with primary antibodies against RGS14 (NeuroMab) and WFS1 (Proteintech) diluted 1:300 (for both) in PBS containing 3% bovine serum albumen (BSA) and 0.2% Triton X-100. Sections were rinsed three times with PBS and incubated with the appropriate secondary antibodies (Invitrogen and Jackson Immunoresearch Laboratories), diluted 1:500, and Hoechst33342 (1:1000; Thermo Fisher Scientific) for 1 h at room temperature. Images were acquired using a TCS SP8 confocal laser-scanning microscope (Leica).

### FC lesion T-maze study

#### Subjects

Male LE rats (n = 22), individually housed with a 12-12 h light-dark cycle, were used in the study. All behavioral experiments were conducted during the light cycle. All protocols were approved by the Institutional Animal Care and Use Committee of Seoul National University.

#### Apparatus

An elevated T-maze (stem, 72 × 8 cm; arms, 40 × 8 cm) containing a food well (2.5 cm diameter, 0.8 cm deep) located at the end of each arm was used(Lee et al., 2018). The food well was covered by a plastic washer to prevent the rat from sampling the reward in the food well from the stem. A quarter piece of cereal (Kellogg’s Froot Loops) was used as reward. The arms of the maze were surrounded by an array of three LCD monitors. A start box with a guillotine door was placed at the end of the stem. Choice latency was measured using infrared sensors installed in front of the start box and in the stem. Behavioral experiments were controlled by a Matlab-based program, and sensor data were acquired using a PCI-6221 data-acquisition device (National Instruments, Austin, TX, USA). Five visual scenes with the following patterns were used in the behavioral experiments: black screen, zebra, pebbles, peacock, and palm tree. Zebra and pebble scenes were presented during pre-training and retrieval sessions, and the reward arm was fixed to one of the arms associated with the scene (i.e., the left arm for the zebra scene and the right arm for the pebbles scene). In the acquisition session, peacock and palm tree scenes were presented across trials (peacock for the left arm and palm trees for the right arm). The scene pairs (zebra-pebble and peacock-palm trees) and their associated reward locations (left and right arms) were pseudo-randomly assigned. The maze was located within a circular curtained area in which white noise was played through loudspeakers. After each session, the maze was vacuumed and wiped with 70% ethanol.

#### Handling and shaping

After 5 d of handling and taming, each rat was habituated to the maze. During habituation, the animal was allowed to freely explore the maze and collect quarter pieces of cereal scattered throughout the maze to become familiar with the maze and its environment. Once familiarized, the rat was subjected to a shaping protocol in which it learned to run directly to a food well and retrieve the reward underneath the covering washer when the start box was opened by an experimenter. In a given session, access to either left or right arm of the maze was blocked by a transparent, heavy acrylic block placed at the entranced of the prohibited arm. Pre-training started after the rat had run a total of 90 trials.

#### Pre-surgical training with old scenes

A training session was composed of sample and test blocks. There were five sample blocks, 10 trials per block. Two scenes (zebra-pebbles or peacock-palm trees) were presented in an alternate fashion across the sample blocks, except between the first and second blocks, when a black screen was presented. Reward arms were identical between first and second sample blocks. For sample blocks, access to the unrewarded arm was blocked by a transparent acrylic block. After the last sample block, the testing block commenced (40 trials) without delay. In the testing block, both arms were accessible from the stem, and one of the two scenes from the sampling block appeared pseudo-randomly across trials. When the rat reached criterion (accuracy for both scenes > 75%), the animal was assigned to either the lesion or control group for surgery.

#### Neurotoxic lesion

Colchicine or sterile saline was injected into the FC for the lesion group (n = 14) and control group (n = 8), respectively. For surgery, the rat was first anesthetized with sodium pentobarbital (Nembutal, 65 mg/kg, I.P.), and its head was fixed in a stereotaxic instrument (Kopf Instruments, Tujunga, CA, USA). Anesthesia was maintained with 1-2% isoflurane (Piramal). Small burr holes were drilled for injection of colchicine (7 mg/mL) or sterilized saline with a microinjection pump (KDS-101; KD Scientific); solutions (0.05 *μ*L for angled injection or 0.1*µ*L for vertical injection) were injected at a rate of 10 *μ*l/h rate through a glass pipette (HSU-2920109; Marienfeld) connected via polyethylene tubing (PE20, Becton Dickinson) to a 10 *μ*l Hamilton syringe. The following coordinates were used for drug injections: 3.4 mm posterior to bregma, ±1.2 mm lateral to the midline, and 4.2 mm ventral from dura at ±20° angle, or 3.5 mm posterior to bregma, 0 mm lateral to the midline, and 3.5 mm from the superior sagittal sinus without an angle. Colchicine is a well-known neurotoxin that selectively ablates granule cells in the DG. Some studies have reported damage outside the DG (e.g., CA1) by colchicine(Ahn and Lee, 2014, Jarrard, 2002), but we found no evidence for volume shrinkage in other areas in the hippocampus, presumably because of the small injection volume. After allowing the rat to recover from surgery for 10 d, we carried out habituation sessions (90 trials) for 4 d to help the rat re-adapt to the experimental situation.

#### Post-surgical test (main task)

Post-surgical tests started 14 d after surgery and followed testing procedures that were identical to those in the pre-surgical training except that no correction was allowed. On days 14 and 18, memory retrieval was tested by presenting old scenes (zebra-pebble or peacock-palm trees). On day 16, memory acquisition was tested using a new pair of scenes that had not been presented during the training period.

#### Histological analysis

Every third section was collected during sectioning for Nissl staining. The extent of the lesion was quantitatively assessed volumetrically by tracing principal cell layers in CA1, CA2, CA3, and DG regions of the hippocampus in photomicrographs (1x magnification) using Photoshop (Adobe). The area of each hippocampal subregion was measured using ImageJ (NIH). The total hippocampal volume was estimated based on the depth (40 *μ*m) of the histological section and the tissue collection frequency (every third section). The tissue range for volumetry was approximately from 2.9 mm to 4.7 mm posterior to bregma. For FC volumetry, photomicrographs taken at 10x magnification were used. Tracking of boundaries of cell layers in the FC in the same manner as used for other cell layers of the hippocampus did not accurately reflect the actual volume of that region owing to the low density of cells with small cell bodies. Therefore, we converted the colored photomicrographs to black-and-white photos and set a threshold at which only the contours of neurons were visible. Measuring the dark areas in the photo afterward allowed fairly accurate volumetric measurement of the FC (**Fig. S4**). Three rats in the lesion group were excluded from analysis for lack of a lesion. Thus, eleven rats were assigned to the lesion group, with eight rats assigned to the control group (**Table S3**).

#### Analysis of behavioral data

Behavioral performances of rats in control and lesion groups were compared by conducting repeated-measures mixed ANOVA and t-test (both paired and unpaired) using Statview (SAS Institute, Cary, NC, USA) or JMP11 (SAS). For post-hoc analyses of ANOVAs, planned comparisons between groups were conducted with Bonferroni correction. Latency was measured from the opening of the start box door to displacement of the washer covering the food well. Trial latency was defined as the time from opening of the start box door to triggering of the last sensor on the track, because hesitation behavior was clearly distinguished in this section. Trial latency was analyzed by Mann-Whitney U test and sign-rank test. Data and statistical results are presented as means ± standard errors of the mean (S.E.M.) or median with individual samples. Statistical results are described in **Table S4**.

### Spontaneous object exploration

#### Subjects

Male LE rats (n = 30), individually housed with a 12-12 h light-dark cycle, were used in the study. All behavioral experiments were conducted during the light cycle. All protocols were approved by the Institutional Animal Care and Use Committee of the Seoul National University.

#### Apparatus

The experiment was carried out in a black square acrylic box (70 × 70 × 60 cm) with a white cue card (40 × 54 cm) attached to the north wall. Environmental noise was masked by playing white noise through loudspeakers throughout the experiment. The floor of the square arena was covered with a large sheet of brown paper to block out local cues on the floor; the paper was replaced between tasks and wiped with 70% alcohol. Magnets were attached below the floor to allow placement of objects in consistent positions across blocks. Experiments were recorded with a digital camera (Logitech HD Pro Webcam C920; Logitech, Newark, CA, USA).

#### Surgery

Rat preparation and surgery for injection of colchicine or saline was performed as described above (*Neurotoxic lesions*). The following coordinates were used for drug injection: 3.5 mm posterior to bregma, 0 mm lateral to the midline, and 3.5 mm ventral from the dura with no angle through the superior sagittal sinus. After surgery, rats were allowed to recover for 6 d. Platform habituation was conducted for 4 d beginning on the next day (day 7).

#### Experimental procedure

On days 7–10, rats were habituated to the platform for 15 min, during which the rat was allowed to freely explore the empty box. Beginning on day 11, a spontaneous object-exploration experiment was conducted every other day under different conditions in the following order: (a) *object location*, (b) *object-in-place (swap)*, and (c) *object-in-place (copy)* (see below). For each task, a different set of objects was used; thus, the same object was never repeatedly used in different tasks.

#### (a) Object location

Two identical objects and copies of the sampled objects were used in the test. The sample block was repeated three times with a 3-min inter-block interval. In a sample block, two identical objects were located in the arena, one in the upper left position and the other in the upper right corner. Rats were allowed to explore the environment for 5 min. Three minutes after the end of the last sample block, the test block was started and one of the objects was displaced to a novel location. The displaced object was pseudo-randomly chosen for different rats. Each test block lasted 5 min. Brown parcel paper covering the floor was changed between blocks.

#### (b) Object-in-Place (swap)

In this task, three different objects were used. The sample block was repeated three times with a 3-min inter-block interval. In a sample block, three identical objects were placed in a certain spatial configuration and rats were allowed to explore the environment for 5 min. A 5-min test block started after the last sample block. In the test block, two objects were pseudo-randomly selected and their locations were swapped. Brown parcel paper covering the floor was changed between blocks.

#### (c) Object-in-Place (copy)

All procedures were the same as in the *object location* test above except that, in the test block, one of the objects was replaced with a copy of the other object.

#### Novel object recognition (5 min or 1 h)

Procedures for the sample blocks were the same as those in the *object location* test above. The test block started either 5 min or 1 h after the last sample block. In the test block, one of the objects was replaced with a novel object.

#### Histological analysis

Histological procedures were the same as those in the lesion study using the T-maze, described above.

#### Analysis of behavioral data

The exploration time for each object was measured manually by the experimenter using EthoVision software (Noldus, Leesburg, VA, USA). Object exploration behavior was registered manually when the direction of the head faced toward an object and the distance between the nose and the object was less than 3 cm. Rearing and biting behaviors were not counted as exploration and were excluded from analyses. For conditions in which three objects were used (i.e., object-in-place and swap), the average time for the two swapped objects was defined as the exploration time for the target object. The discrimination index was defined by subtracting the target object-exploration time from the familiar object-exploration time divided by the sum of the exploration time for both objects. For statistical analyses, the Wilcoxon signed-rank test was used for comparing exploration times between objects, the Mann-Whitney rank-sum test for comparing between groups, and the one-sample sign test for comparison against the chance level. Statistical analyses were conducted using Statview (SAS Institute) and JMP11 (SAS Institute). Median and interquartile ranges were used for presentation of data and statistical results. Statistical results are described in **Table S5**.

### Electrophysiological experiment (random foraging)

#### Subjects

Male LE rats (n = 12), individually housed with a 12-12 h light-dark cycle, were used. All behavioral experiments were conducted during the light cycle. All protocols were approved by the Institutional Animal Care and Use Committee of the Seoul National University.

#### Apparatus

The experiment was carried out in a black, square acrylic box (70 × 70 × 60 cm) with a white cue card (40 × 54 cm) attached to the north wall. Environmental noise was masked by playing white noise through loudspeakers throughout the experiment. The floor of the square arena was covered with a large sheet of brown paper to block out local cues on the floor; the paper was replaced between tasks and wiped with 70% alcohol. A digital camera, commutator (PSR-36; Neuralynx), and a custom semi-automatic feeder were installed in the ceiling. The feeder scatters chocolate sprinkles when an experimenter pushes a switch from outside the testing room while monitoring the rat’s movement.

#### Hyperdrive implantation

Detailed procedures for constructing and surgically implanting the hyperdrive can be found in our previous studies(Lee et al., 2018). Briefly, a custom-made hyperdrive containing 24 nichrome wire tetrodes (17.8 *μ*m diameter; A-M Systems, Sequim, WA, USA) was implanted in rats (11–41 wk old; n = 12). To avoid damage to the superior sagittal sinus, we angled the tetrode-carrying bundle of the hyperdrive at 20° to approach the FC obliquely. The hyperdrive itself was also inserted into the brain at an angle (10°). The surgical coordinates were pre-calculated to target the right hemisphere of the FC with the tip of the tetrode bundle (9 G or 12 G stainless-steel tubing), aligned 3.6 mm posterior to bregma and 2.2 mm lateral from the midline. A stainless-steel wire coupled to the ground channel of the hyperdrive was connected to a skull screw over the cerebellum for use as an animal ground.

#### Electrophysiological recording

After 7 d of recovery from surgery, tetrodes were lowered daily into the target areas (FC and CA1) over the course of approximately 3 wk. Spiking activities from single units and local field potentials (LFPs) were fed to the data-acquisition system (Digitalynx SX; Neuralynx) through an electrode interface board (EIB-36-24TT; Neuralynx), pre-amplifier (HS-36; Neuralynx), and tether (HS-36 Litz tether; Neuralynx). Signals were digitized at 32 kHz, filtered at 600-6,000 Hz, and amplified 1,000-10,000 times. Neural signals were recorded during sleep in a sound-attenuating booth outside the experimental room before (pre-sleep) and after (post-sleep) each behavioral recording session. The rat was allowed to sleep in the booth for at least 30 min to obtain sufficient resting period-associated neural data. LFP data relative to animal ground was also recorded with simultaneous recording of single units relative to the reference wire of rats (n = 10). Only LFP data against the animal ground was used for LFP analysis.

#### Post-surgical behavior

After most of the tetrodes had reached their target areas, a habituation session was started in the foraging box with the tether connected to the hyperdrive at least for 3 d, followed by random foraging sessions for 6-14 d. During the foraging session (15 min), rats were allowed to move freely around in the square box and consume randomly scattered chocolate sprinkles while hippocampal neural signals were recorded.

#### Histological verification of electrode positions

After the last recording session, the rat was sacrificed with an overdose of CO_2_ and transcardially perfused first with PBS and then with a 4% formalin solution. The head was fixed in formalin solution for 1 d before the brain was extracted. The extracted brain was fixed in a 4% formaldehyde solution containing 30% sucrose at 4°C until it sank to the bottom of the tube. The brain was then coated with gelatin and fixed in sucrose-formaldehyde solution for 1-2 d. The brain was sectioned at 40 *μ*m using a freezing microtome (HM 430; Thermo-Fisher Scientific), and every section was mounted on a glass slide and thionin stained. Photomicrographs were taken with a digital camera (DS-Fi1c; Nikon) attached to a fluorescence microscope (Eclipse 80i; Nikon), and tetrode tracks were reconstructed three-dimensionally (Voxwin, UK) to match between the electrode track with the pre-configured bundle design. The recorded depth of tetrodes and physiological profiles were also considered during reconstruction procedures. Tetrodes anterior to a specific point (3.0 mm from bregma) or located in the border between subregions were excluded from the analysis.

#### Unit isolation

Single units were recorded simultaneously from the FC (n = 228) and CA1 (n = 1195). Units were isolated manually using custom software (WinClust), with the peaks of waveforms used as the major measurements, as described previously(Lee et al., 2018), while also considering other parameters (e.g., energy). Spikes collected during sleep sessions were also used during unit isolation for judging the stability of recordings. Only single units that meet the following criteria were included in the analysis: (a) > 50 spikes identified in a cluster in both pre-sleep and post-sleep sessions, and < 1% of total spikes occurring within the refractory period (1 ms); (b) mean firing rate of a cluster < 10 Hz and only putative complex spiking neurons included; and (c) units are ‘place cells’ (i.e., the number of spikes in the open field ≥ 50, spatial information ≥ 0.5 with a *p*-value < 0.05; see ‘Electrophysiological data analysis’ for details). Spikes recorded when the animal was relatively stationary (moment velocity < 5 cm/s) were filtered out and were not used in the analysis(Henriksen et al., 2010). For delineating physiological boundaries, CA1 was divided into two subdivisions (dmCA1 and dCA1) based on an online atlas(Boccara et al., 2015): (1) the distalmost part of the CA1 (dmCA1), which was considered the FC in some previous reports(Boccara et al., 2015, Henriksen et al., 2010), and (2) the distal CA1 (dCA1), which is located laterally relative to the dmCA1. Only tetrodes located 3 mm posteriorly from bregma were used, and tetrodes located at the border between subregions were excluded from the analysis. After applying all criteria, the final number of place cells analyzed was as follows: FC, n = 40; dmCA1, n = 170; and dCA1, n = 104.

#### Electrophysiological data analysis

Spike width was defined as the temporal distance between the peak and trough of the averaged spike waveform. The averaged spike waveform was constructed using all channels and spikes during a recording, including sleep sessions and behavior sessions. A burst index was defined as the number of spikes in 3-5 ms bins divided by the average number of spikes in 200-300 ms bins(Senzai and Buzsaki, 2017). Position data from the foraging session was binned into a matrix (bin size = 2.7 × 2.7 cm^2^) to construct a rate map, and spike data were assigned to the matching bins. A rate map was calculated by dividing the number of spikes by the duration of visit per bin (firing rate). The resulting rate map was smoothed using an adaptive binning method(Skaggs et al., 1996).

In the rate map of a unit, the unit’s place field was defined as continuous bins showing > 20% of the peak firing rate. At least 20 continuous bins (88.2 cm^2^) were required for a unit’s firing field to be qualified as a place field. In-field firing rate was measured as the firing rate within the place field. Spiking activity spatial information was measured as follows(Skaggs et al., 1993):

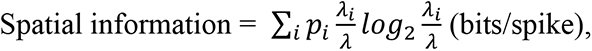

where *i* denotes the bin number, *p_i_* and *λ_i_* represent the occupancy rate and firing rate of the *i*^th^ bin, respectively, and λ denotes the mean firing rate. The probability of obtaining a spatial information score was calculated from the random distribution of spatial information using a Monte Carlo method. Specifically, the spiking train of a unit was randomly shifted (minimum shifting unit = 33 s), and spatial information was re-calculated from the generated rate map for each shift. This procedure was repeated 1000 times. The P-value was defined as the proportion of shuffled data above the spatial information obtained from the actual rate map.

Coherence was measured as the correlation coefficient between the raw rate map and the reconstructed rate map, where the firing rate of each pixel was the averaged firing rate of adjacent pixels. The stability of a rate map was measured as the correlation coefficient (R) between the rate maps constructed for the first half and second half of a session (each half = 7.5 min) in the raw rate map. The firing rate (sleep) was measured during sleep sessions after the behavioral experiment. The commercial software packages JMP (SAS), Statview (SAS), and Matlab (MathWorks, Natick, MA, USA) were used for statistical comparisons (ANOVA, repeated-measures ANOVA, t-test, paired t-test, and one-sample t-test). Statistical results are described in **Table S6**.

#### LFP analysis

The analysis procedures followed those in our previous report (Ahn et al., 2019) and used a custom Matlab (MathWorks) program. Briefly, after the LFP data were down-sampled (1/16) to 2 kHz, LFP data in the 150–250 Hz range were filtered with a bandpass filter. The parameters for the filtering were as follows: passband edge frequency (Wp), 145–255 Hz; stopband edge frequency (Rp), 130–270 Hz; passband ripple (Rp), 3dB; stopband attenuation (Rs), 15 dB; filter type (ftype), bandpass. These parameters were used for Matlab functions in the signal processing toolbox (MathWorks) in the order, ‘buttord’, ‘butter’ and ‘zp2sos’. The filtered EEG was derived from the ‘filtfilt’ function.

#### Ripple detection

For ripple detection, filtered LFP data were smoothed using a Gaussian filter, and envelopes were detected. For calculating the envelope, the ‘smoothdata’ function was used after a Hilbert transform of LFP data (smoothing bin size: 25/2000 s). The envelope was defined as LFP data > 1.5-times the standard deviation (STD) from the mean. For noise reduction, data points > 11 STD were discarded from analyses. Also, ripples less than 20 ms were discarded, and ripples with intervals < 20-ms were considered a single ripple. Once the ripple boundary for each tetrode was confirmed, the border was extended to one STD from the average of the envelope. The ripple of a session was defined as the total duration of overlapped ripples recorded in CA1 (both dCA1 and dmCA1), and ripples detected by at least three tetrodes were included in the analysis to reduce noise.

#### Ripple power analysis

Once ripple detection was complete, ripple power was measured in unfiltered LFP data (unit: mV) using the ‘mtspectrumc’ function in the Chronux 2.0 toolbox(Bokil et al., 2010) with the following parameters: frequency (Fs), 2000 Hz; time band width and multiple tapers (tapers), [3 5] (300 ms); frequency to be analyzed (fpass), 300 Hz; Pad (pad), 0; Trial (trialave, not used), 1; mean and S.T.E. (err, not used), [1 0.05]. The power in a ripple was measured by the ‘bandpower’ function in Chronux. Because LFP data were measured against the animal ground, body movements affected LFP power. We noticed that the corpus callosum (CC) showed ripple power similar to CA1 when the rat was in an awake state while showing less ripple power than CA1 in the sleep state. Therefore, we rejected ripples whose power ratio of averaged CA1 and averaged CC was less than 1.5. Power was transformed into decibel (dB) using the ‘pow2db’ function in Matlab(Rasmussen et al., 2017). The outburst ratio was defined as the average firing rate in the boundaries of ripples divided by the average firing rate out of ripples. Only LFP data recorded from the post-experimental sleep session was used. The representative peri-event raster plot and histogram were constructed using NeuroExplorer (Nex Technologies, Vienna, Austria). Statistical results are described in **Table S7**)

### Electrophysiological experiment (visual contextual task in the T-maze)

Most electrophysiology procedures, including surgery protocol, electrophysiology recording procedure, histology and analysis, were the same as those used for the random foraging experiment. Only unique procedures are described here.

#### Subjects

Male LE rats (n = 4), individually housed with a 12-12 h light-dark cycle, were used in the study. All behavioral experiments were conducted during the light cycle. All protocols were approved by the Institutional Animal Care and Use Committee of the Seoul National University.

#### Apparatus

An elevated T-maze (stem, 73 × 8 cm; arms, 40 × 8 cm) was used. The experiment setup is similar to that described in the lesion study/T-maze section, but this experiment was conducted in another room. A food well (2.5 cm diameter, 0.8 cm deep) was located at the end of each arm. A quarter piece of Froot Loops (Kellogg’s) was used as a reward. The arms of the maze were surrounded by an array of three LCD monitors. A start box with a guillotine door was placed at the end of the stem. Latency was measured using infrared sensors installed in front of the start box and in the stem. The behavioral experiment was controlled using a Matlab-based custom program, and sensor data were acquired using a data acquisition device (PCI-6221; National Instruments). Seven visual scenes were used in the behavioral experiments: blank screen for habituation and first sample block; zebra and pebbles scenes for retrieval; and bamboo, mountain, peacock, and palm tree scenes for acquisition. A digital camera and commutator (PSR-36; Neuralynx) were installed in the ceiling.

#### Behavioral procedure

##### Pre-surgical training

The pre-surgical training procedures were identical to those described for the lesion study using the T-maze.

##### Post-surgical training

After the recovery period (1 wk), the rat was subjected to block habituation sessions until the tetrodes reached the target cell layer in the FC and CA1. No scenes were displayed to the rat during this period except the black screen. The reward location was changed between sessions but not within a session. When most tetrodes were prepared to record single units from the target regions in the hippocampus, the main recording started (see below).

##### Post-surgical test

Testing for retrieval and acquisition and spatial shuttling was conducted for 6 d using physiological recordings. The number of trials in the test block was extended (80-120 trials) compared with the lesion study to acquire as much neural data as possible. Retrieval and acquisition sessions were performed in an alternate fashion for 4 d. For the retrieval session, zebra and pebble scenes were presented. Only one session was performed per day. For the first acquisition session, peacock and palm tree scenes were presented, whereas bamboo and mountain scenes were presented for the second acquisition session. If the rat showed performance less than 70% during the test block, data from the session was discarded and the session was repeated the next day. This constraint resulted in two acquisition sessions from two rats being discarded and repeated. Two shuttling sessions were conducted after the tasks. During a shuttling session, no scene was presented, and the reward location was fixed. Reward locations were changed between sessions.

#### Unit isolation

All procedures were the same as those described for the foraging experiment. Briefly, to include only putative complex spiking neurons, we excluded single units that did not meet the following criteria from the analysis: (a) >50 spikes identified in a cluster in both pre-sleep and post-sleep sessions; (b) <1% of total spikes occurring within the refractory period (1 ms); and (c) mean firing rate of a cluster < 10 Hz (dmCA1, n = 427; FC, n = 112). Only place cells at the outbound trajectory were included (number of spikes ≥ 50, spatial information ≥ 0.5, spatial information p-value < 0.05; dmCA1, n = 257; FC, n = 52). Also, only place cells for which place fields represented the stem (>20% of peak firing rate and >4 consecutive bins) and with an episodic field (see below) were included in the analysis. The final number of place cells analyzed was as follows: FC, n = 10, 14 and 8 in the acquisition task, retrieval task, and shuttling session, respectively; dmCA1, n = 58, 50, and 40 for the acquisition task, retrieval task, and shuttling session, respectively.

#### Electrophysiological data analysis

Common procedures overlapping with those used for the foraging experiment are not repeated here. Briefly, position data from behavior sessions were binned into a matrix (bin size = 10 × 10 px^2^ = 2 × 2 cm^2^) to construct a rate map. The place field was defined as bins where firing rate was more than 20% that of the peak firing rate. If the length of a place field was less than 4 bins (8 cm) or had no episodic fields, the place field was excluded from the analysis. A rate modulation index was defined as the difference in in-field firing rates between scene conditions divided by the sum of in-field firing rates.

The raw spike map and rate map were constructed to show the time course of place cell activity in the FC. Maps were constructed only for the stem in which both trajectories in each condition overlapped. The rate map was constructed using the adaptive binning method, and each bin along the abscissa was composed of five trials (bin size = 2 cm × 5 trials). The trial-position map of each place field was averaged and linearized to present the firing rates of each trial bin. An episodic place field was defined as clusters with firing rates greater than 20% of the peak firing rate in the linearized map. Episodic place fields spanning less than four bins (20 trials) were excluded from the analysis. Also, place fields without a spike in more than 70% of trials were excluded. The transient duration of place cell activity was defined as the temporal distance from the beginning of the start trial to the last trial.

A population matrix that described the distribution of place fields across trials was constructed using Boolean coding. If a place field was present in a specific trial bin (1 bin = 5 trials), the bin was assigned a value of 1; for simplification, other trial bins were coded as zero regardless of the position of the place field, scene, correctness, or task. The network state was defined as a Boolean vector of the trial bins coding the on- and off-state of the place cell population. The distance between network states was measured as the Euclidean distance between two adjacent trial bins in an individual vector space whose main axes were individual cells. Because the number of place cells in the dmCA1 was larger than that in the FC, cells in the dmCA1 were randomly subsampled 1000 times so that each subsample size matched the sample size of the FC. For visualization purposes, the dimension of individual space was reduced using the PCA function in Matlab (MathWorks). However, this restriction was only applied for visualization and not for data analysis. The cumulative distance corresponds to the sum of the distances from the first distance bin to a specific distance bin. The cumulative distance up to 130 trials and Monte-Carlo shuffling were used for statistical comparisons. The P-value was defined as the number of subsamples in the dmCA1 that exceeded the cumulative distance of the FC. Recording stability was defined as the difference in firing rates between pre-sleep and post-sleep sessions divided by the sum of the two. Statistical results are described in **Tables S8** and **S9**.

## Supplementary Materials

**Figure S1.**
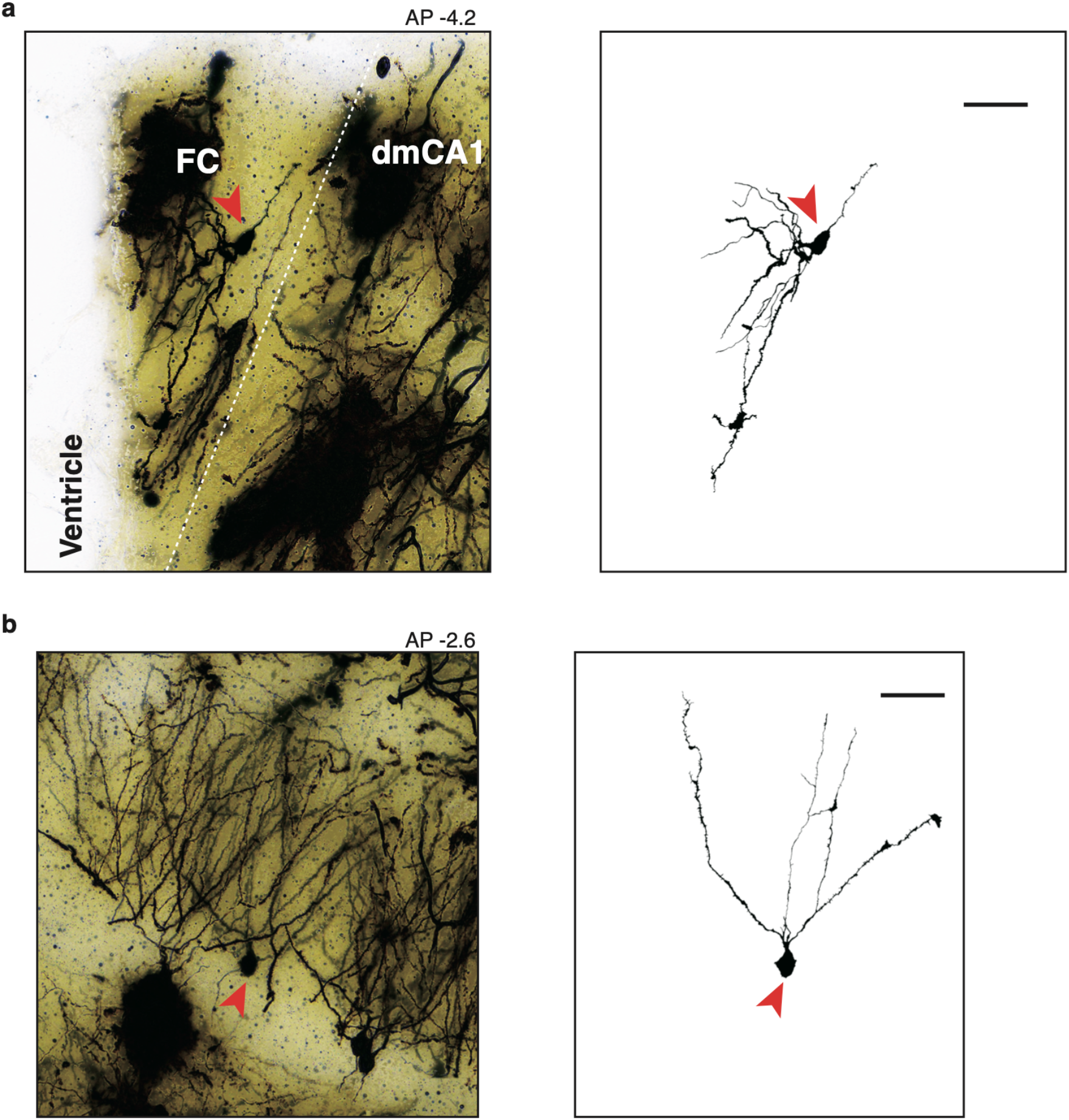
Granule cell comparison between the FC and DG. **a.** A granule cell in the FC. A Golgi-Cox-stained FC cell (left) and single-cell morphology (right) are shown. The red arrowheads indicate the same neuron. A single cell in the FC shows cone-shaped dendrites and the dendritic branches directed toward the external surface (ventricular surface), common features of granule cells in the DG ^14^. However, the overall dendrite shape seemed to be distorted by the shape of the FC. Scale bar: 50 *μ*m. **b**. A Golgi-Cox-stained section including DG granule cell (left) and single granule cell morphology (right) (red arrowhead). As previously described, the DG granule cell showed cone-shaped dendritic arborization and the dendritic branches directed toward the external surface^14^. Scale bar: 50 *μ*m.

**Figure S2.**
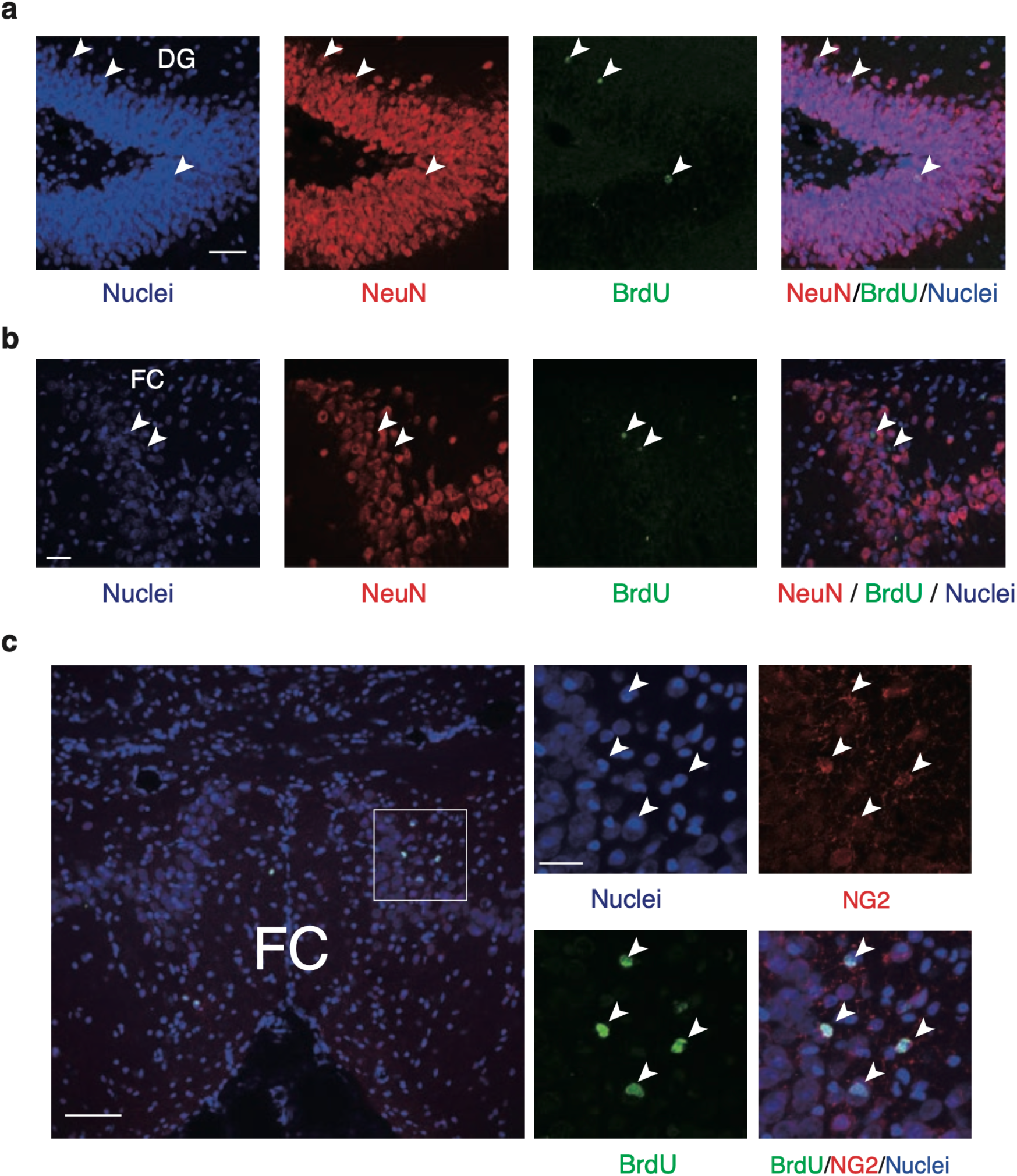
Absence of neurogenesis in the FC. **a.** Some granule cells (NeuN^+^, red) in the DG are newborn cells (BrdU^+^, green; arrowheads) as marked by the arrowheads. The merged image of the two photomicrographs (NeuN + BrdU) overlapped with the nuclei-stained section (Hoechst staining, blue) showed the presence of adult-born granule cells in the DG (white; arrowheads). Scale bar: 50 *μ*m. **b.** In the FC, the locations of BrdU^+^ cells (arrowheads) did not overlap with the locations of the NeuN^+^ cells. Scale bar: 30 *μ*m. **c.** BrdU-labeled cells in the FC were co-localized with the NG2-labeled glial cells (oligodendroblasts; arrowheads). Scale bars: 75 *μ*m (*left*) and 25 *μ*m (*right*).

**Figure S3.**
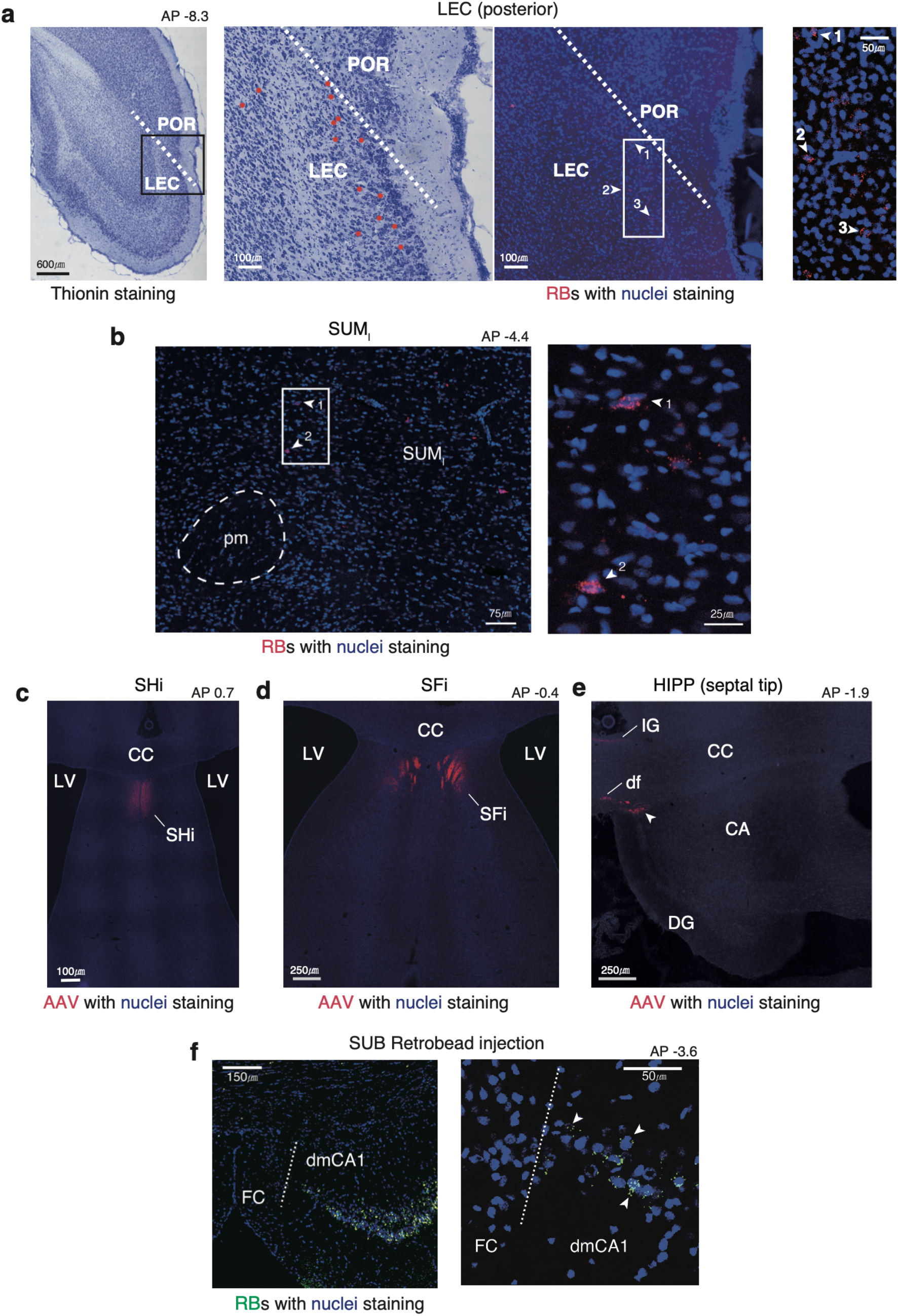
Afferent and efferent connections of the FC. **a.** Afferent projections of the FC coming from the posterior LEC. The leftmost photomicrograph shows the overall cortical structure and the boundaries between the LEC and POR (white dotted line). The second one is the enlarged picture of the box located in the first picture. The third picture from the left shows the adjacent, fluorescent-stained section. Red dots denote the locations of retrobeads-positive cells shown on the fluorescent section. The last picture is the enlarged view of the white box in the third picture. **b.** Retrobeads injected into the FC were detected in the lateral part of the supramammillary body. The projection was supported by SUM anterograde tracing study^15^. **c-e**. Efferent connections of the FC to the SHi, SFi, IG, df, and septal tip of the hippocampus (white arrow heads). The septal tip region is also named SFi in the atlas^16^. **f.** *(left)* Afferent projections of the dmCA1 from the subiculum. The green retrobeads were located in the CA1 but not in the FC, showing the distinct anatomical borders between the CA1 and FC*. (center and right)* A magnified view of the border between the FC and dmCA1. Abbreviations: POR = postrhinal cortex, LEC = lateral entorhinal cortex, SUM_l_ = supramammillary nucleus (lateral parts), pm = principal mammillary tract, SHi = septohippocampal nucleus, SFi = septofimbrial nucleus, LV = lateral ventricle, CC = corpus callosum, IG = indusium griseum, df = dorsal fornix, CA = cornus ammonis, DG = dentate gyrus, SUB = subiculum, dmCA1 = distalmostCA1

**Figure S4.**
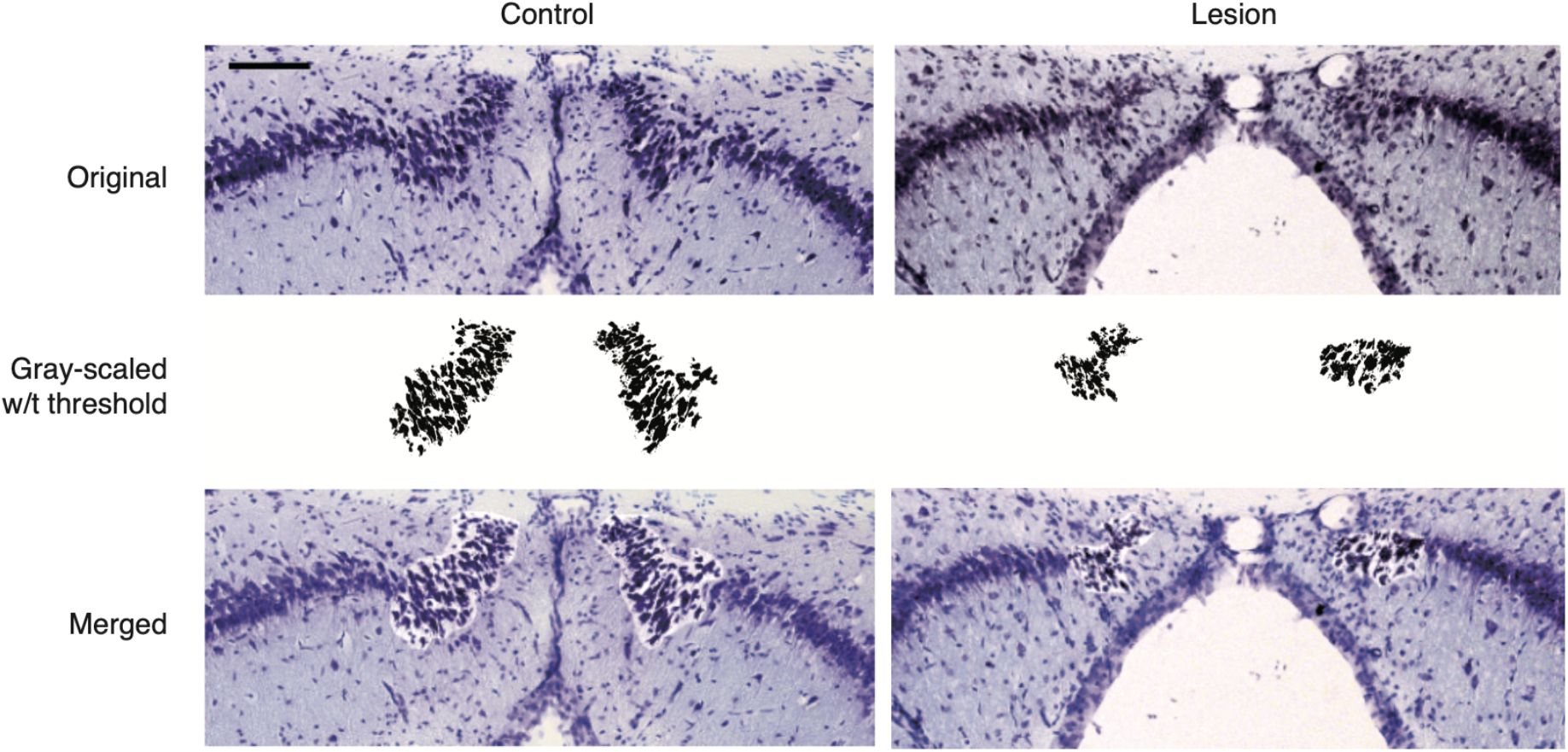
Volumetric measurement of the FC. Original photomicrographs taken with 10x magnification are shown in the first row (Original). Clusters of FC cells isolated from the background by setting an intensity threshold in the black-and-white photos of the original photomicrographs. The bottom row shows the merged photo to verify the locations of the FC in the original photomicrographs. Scale bar = 100 *μ*m.

**Figure S5.**
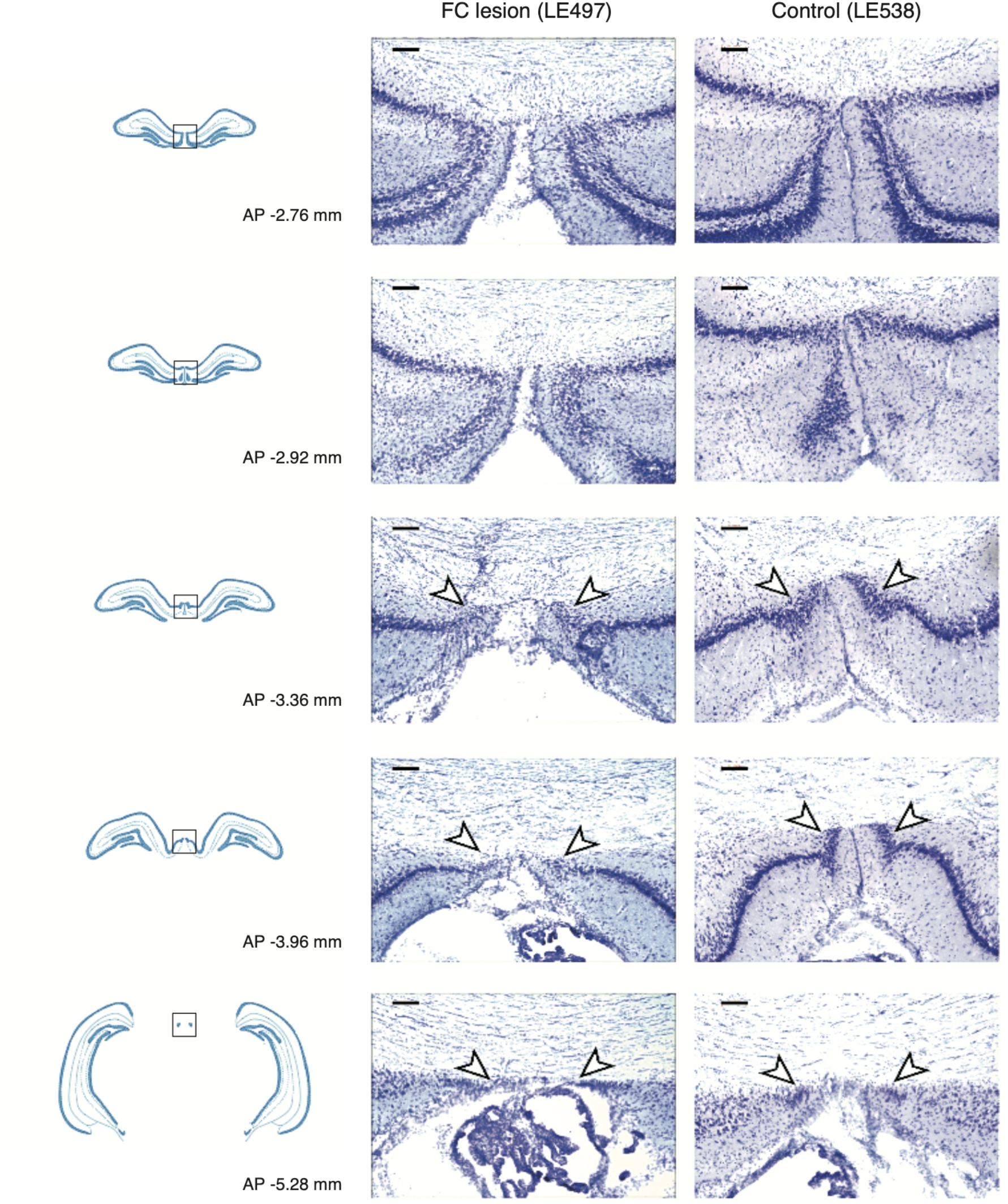
A selective lesion in FC after colchicine injection. The left column shows the hippocampus diagram and distance posterior to the bregma. The next histological pictures show the center of the hippocampus of lesion group (*center*) and control group (*right*) at the corresponding anterior-posterior coordinate. White arrow heads indicate the FC region. The diagram is modified from the atlas of Paxinos and Watson (2009). The scale bar denotes 0.1 mm.

**Figure S6.**
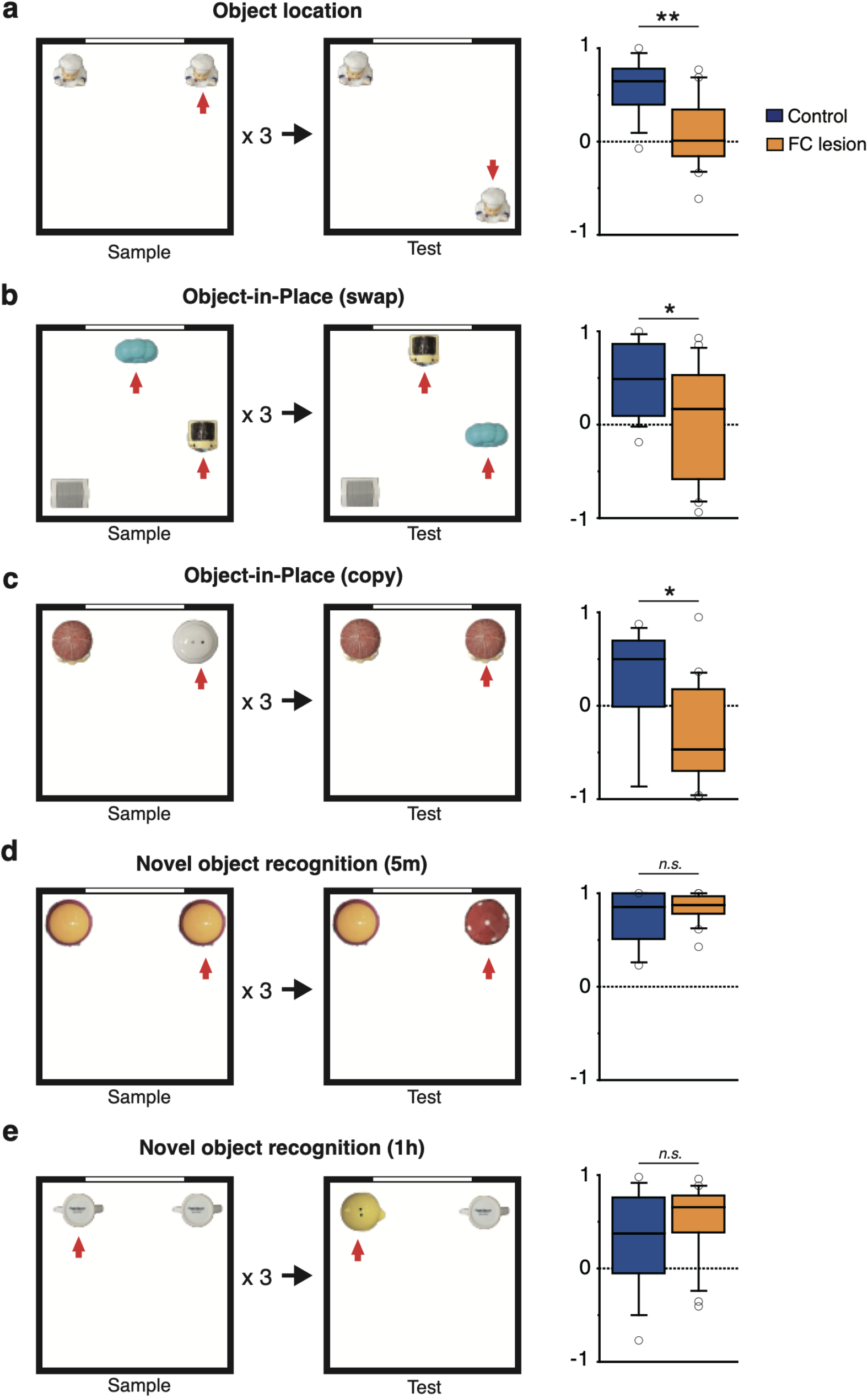
FC lesions impair object-place paired associative memory, but not simple object recognition. The behavioral results from the VCM task suggest that the FC might be critical for setting up a new contextual representation in the hippocampus via the DG network, but not necessarily for retrieving an old one. If that is the case, the FC should be also important for recognizing object-place (or object-context) associative representations, but not when place or context is not critically involved (e.g., simple object recognition). To test this hypothesis, we ran a series of spontaneous object-exploration experiments that required recognizing places, objects in place, and novel objects. These (**a**-**c**) manipulations are known to recruit the DG in the hippocampus ^17-21^. **a.** The FC is important for recognizing a displaced object. when one of the two identical objects was displaced to a new location (red arrow) after being repeatedly presented in fixed locations, the control group preferred to explore the displaced object more than the other object but the FC-lesion group failed to do so (Z = 2.99, P = 0.003). The results suggest that the FC is important for the contextual recognition of objects. The box plot shows the index for the amount of exploration for the displaced object. **b-c.** The FC is important for object-in-place paired associative memory. **b**. Inspired by prior studies ^19-21^, we tested if FC-lesioned rats were able to recognize new object-place paired associates (*object-in-place*) when two of the three objects swapped their locations (*swap*). In the *swap* condition, the rat sampled three different objects for three blocks and, before testing began, two of the objects (red arrows) switched their locations. The box plot shows the index for the amount of exploration for the changed object. **c**. In the *copy* condition, two different objects were explored and one of the objects was replaced with a copy of the other object (red arrow). The box plot shows the index for the amount of exploration for the changed object. In both object-in-place tests, although the objects were all familiar during the test, the FC lesion group was significantly impaired in exploring the object that was encountered in a novel context compared to controls (Z = 1.99, P = 0.047 for the swap test; Z = 2.11, P = 0.035 for the copy test). Our findings suggest that FC may play key roles in setting up discrete contextual representations in the hippocampus with which individual objects can be associated for object-in-place memories. **d-e**. The FC is not important for simple object recognition. To test a possibility that the FC is important for any type of novelty, rats were tested in a simple object recognition task. Prior studies have shown that the hippocampus (including DG) is not important for detecting object novelty alone in the absence of a spatial change ^17,18^. **d**. In this test, rats were repeatedly exposed to two identical objects occupying the opposite corners of a square environment. Then, one of the objects was replaced by a novel object (red arrow). The box plot shows the index for the amount of exploration for the changed object. The amount of the targeted exploration of the changed object in the FC-lesion group was not significantly different from that of the control group, suggesting that FC is not important for detecting a novel object (Z = 0.65, P = 0.517). **e**. A previous study suggested that the hippocampus might play an important role in simple object recognition with a significantly long delay (e.g., 1 hour) between the sample and test phases ^22^. The experiment is the same as (**d**) except the interval between last sample block and testing (1h). The box plot shows the index for the amount of exploration for the changed object. However, the FC-lesion group’s object preference was not significantly different from controls in our study even when we put 1-hour delay before testing (Z = 1.03, P = 0.305). Taken together, the results from our series of experiments strongly suggest that the FC is critical for acquiring unique contextual object representations in the hippocampus presumably through DG. (*p < 0.05; **p < 0.01). Statistical results are described in Supplementary Table 5.

**Figure S7.**
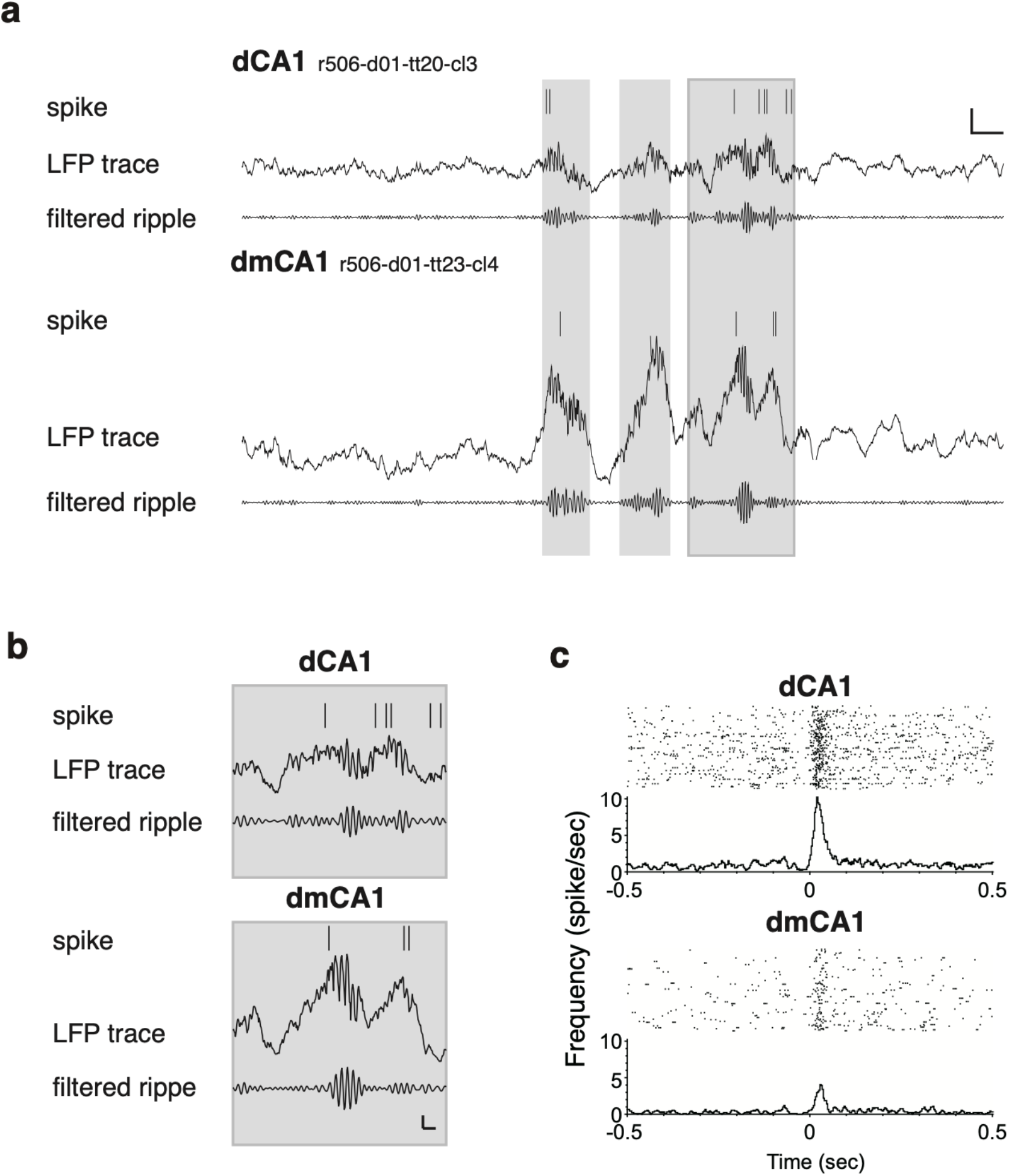
LFP and cellular response of the dCA1 and dmCA1 during the SWR. **a**. Spike train and LFP trace for dCA1 and dmCA1 units during the sleep session. The raw LFP trace and band-pass filtered one (range: 150-250 Hz) are presented. The gray zones exist during the SWR period. The vertical and horizontal scale bars denote 400 *μ*V and 50 ms. **b**. The enlarged figure of the last SWR event in (b). The vertical and horizontal scale bars denote 200 *μ*v and 10 ms. **c**. A peri-event time histogram (PETH) and raster plot (PETR) aligned to SWR onset timing for the representative cells presented in (b) are displayed.

**Figure S8.**
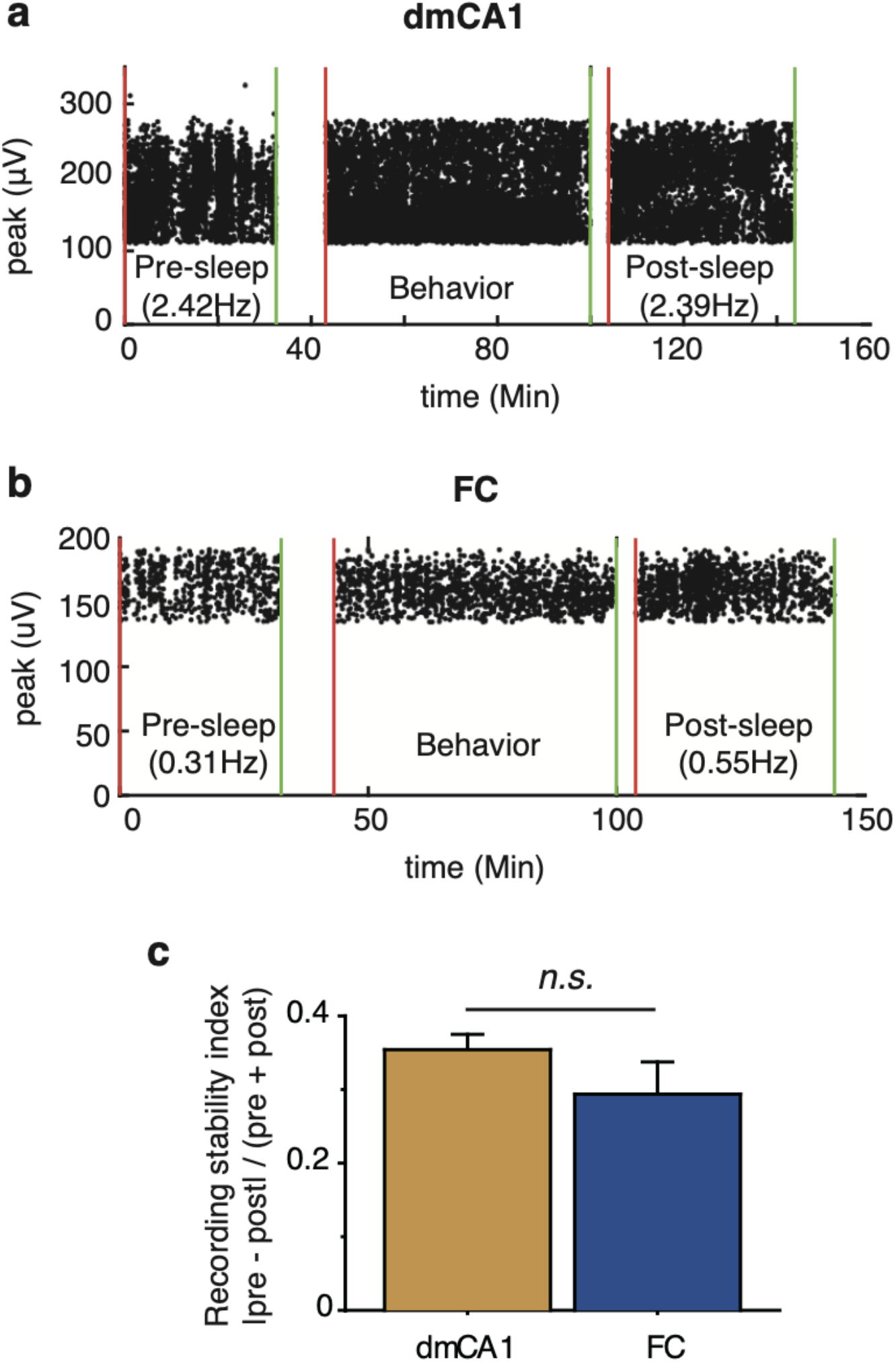
Representative examples of place cell stability. The peak amplitudes of place cells in the dmCA1 (**a**) and FC (**b**) during the sleep sessions and behavior session are displayed. The x-axis indicates the time from the start of the pre-sleep session. **c**. The firing-rate stability index, which is the difference in firing rates between the pre-sleep and post-sleep sessions divided by the sum of the firing rates in both dmCA1 and FC. Statistical results are described in Supplementary Table 8.

**Figure S9.**
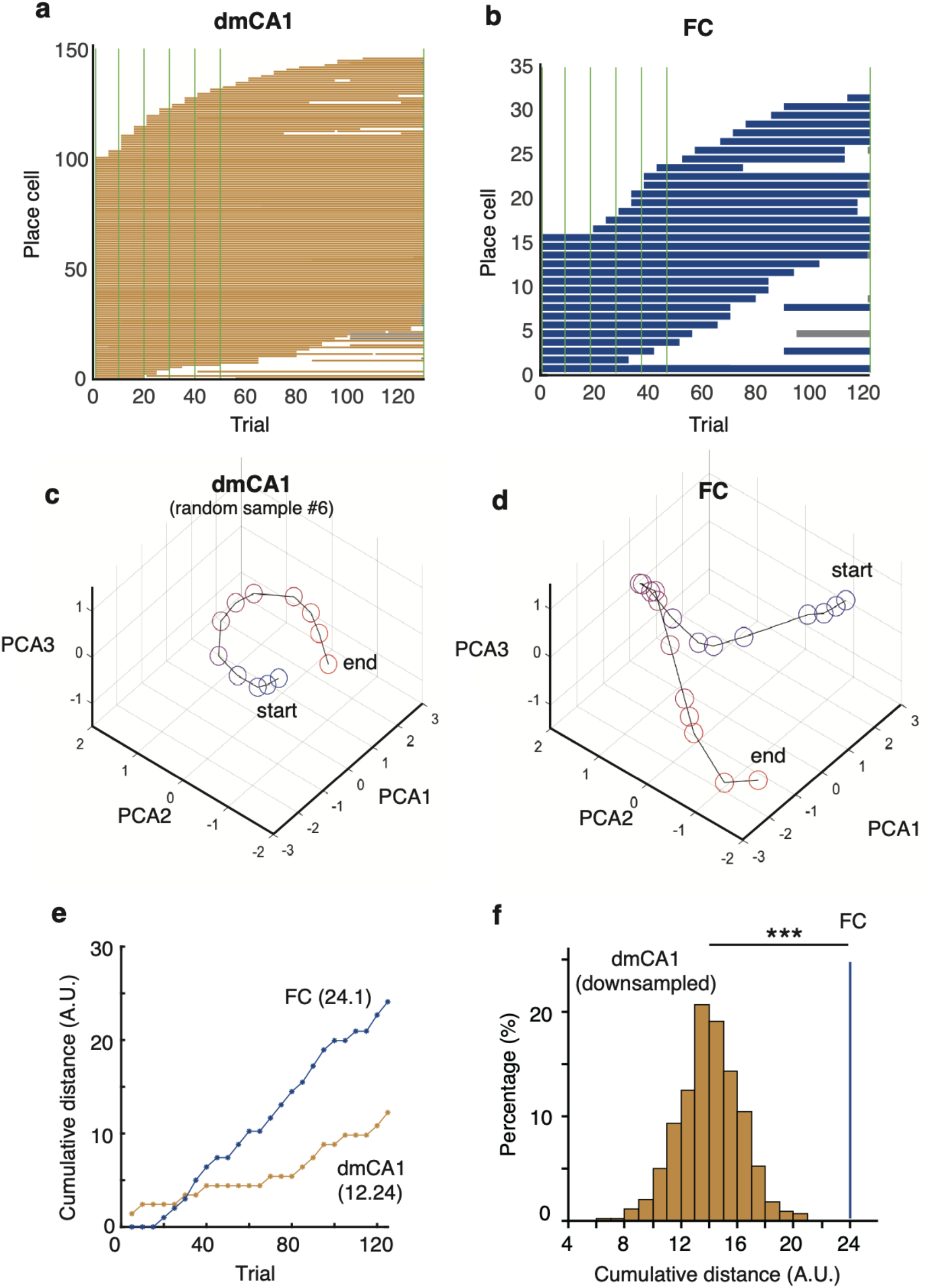
Network state at the population level changes more dynamically over time in the FC than in CA1. **a, b**. Transient spatial coding of place cells in both the dmCA1 (**a**) and FC (**b**). Each place cell’s spiking state (ordinate) was plotted as binary information (0 for no spiking and 1 for spiking) across trials (abscissa). Gray zone indicates that the recording session ended prematurely. Note that spatial firing covers the entire trial of an overall session in most CA1 cells but not FC cells. **c, d**. Representative examples of network state changes (visualized in PCA axes) in the dmCA1 (random sample #6 out of 1000 down-sampled cases; **c**) and the FC (**d**). Blue and red circles indicate first and last trials, respectively. The distance between adjacent trials indicates the abruptness of the change in network state between trials. **e**. Cumulative distance of network states across trials. The example for the dmCA1 is the same down-sampled example shown in (**c**). **f**. The distribution of down-sampled cumulative distances from the starting trial to the last trial in the dmCA1 (orange) and the location of the FC value in the distribution (blue line). Note that the distance traveled in the state space by the FC network was far greater than the highest distance in the CA1 distribution. The cumulative state distances (calculated using all dimensions) across trials for the dmCA1 and FC differed significantly between the two subregions (FC = 24.1, dmCA1 = 14.1 ± 0.1, mean ± S.T.E; ***P < 0.001; Monte-Carlo method).

**Figure S10.**
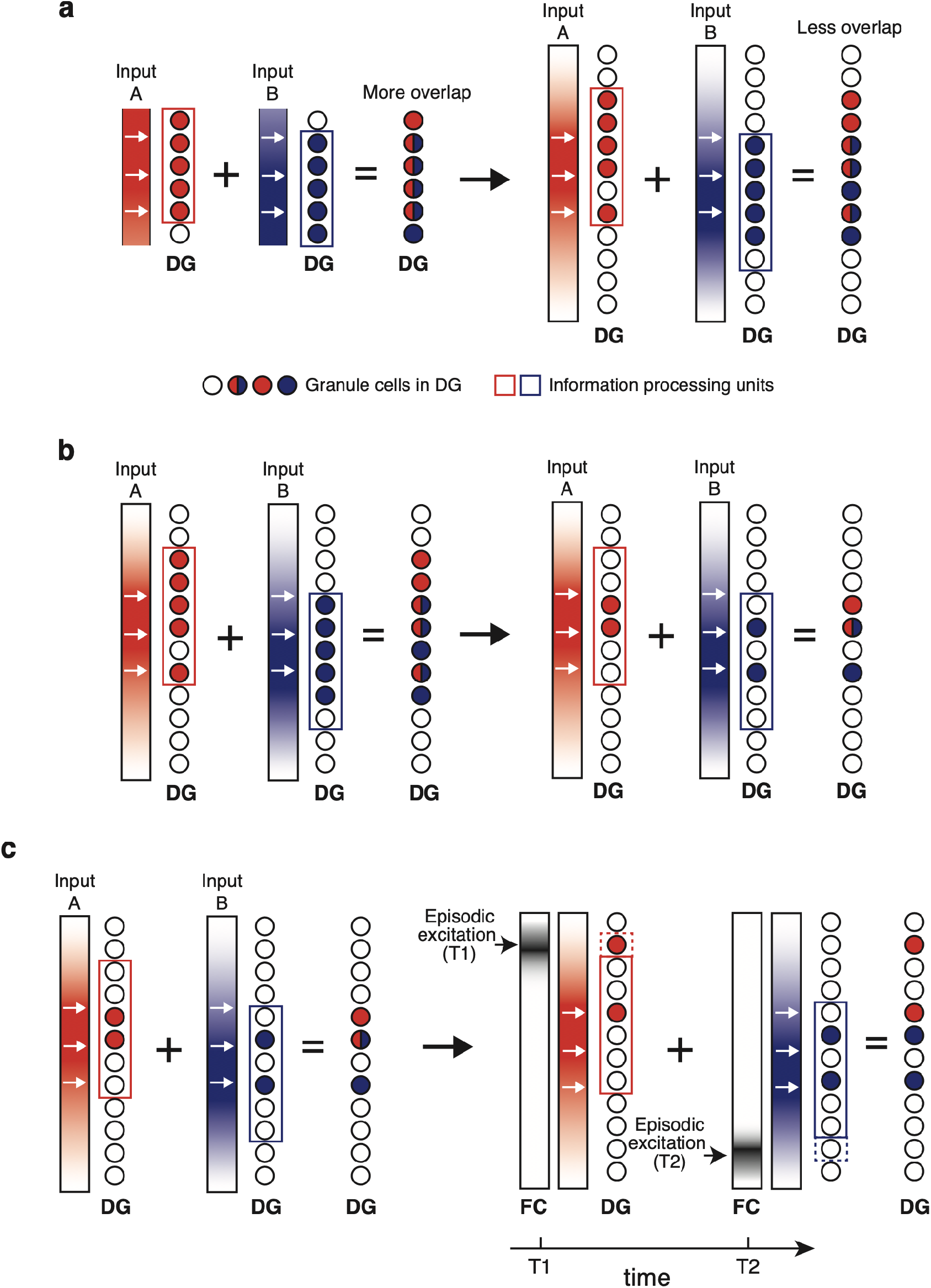
A possible mechanism for the fasciola cinereum to contribute associative learning. **a.** The vertical bars indicate input to granule cells in the dentate gyrus (DG GC), while circles indicate granule cells. The intensive color indicates the strong input to corresponding DG GCs, and red and blue cells indicate representations for the input A and B, separately. DG GC with both colors indicate representation overlap. The colored squares indicate the cell group that has the potential to be activated (‘candidates’). The schematic figure shows how the large capacity reduces the overlap of representations. **b**. The presentation scheme is the same as in (a), except that the number of cells required to form a representation in the DG is reduced. **c**. The presentation scheme is similar to (b), but additional episodic excitation is introduced. The black-and-white vertical bars indicate additional episodic excitation from the FC (extrinsic excitation) or newborn granule cell (intrinsic excitation). The dotted squares indicate an extended candidate due to additional episodic excitation at the point in time T1 and T2.

**Table S1.** Genes known to be expressed in FC and CA2 subregions based on the literature. Genes expressed throughout hippocampal subregions, such as CamKII (Ca^2+^/calmodulin-dependent protein kinase II) and GAD67 (glutamate decarboxylase 67) were excluded.

| Gene | Full name | Reference | Note |
| --- | --- | --- | --- |
| **Genes expressed in the both FC and CA2** |  |  |  |
| Pcp4 | Purkinje cell protein 4 | Lein et al. <sup>23</sup> | Also observed in DG |
| Amigo2 | Adhesion molecule with Ig-like domain 2 | Laeremans et al. <sup>24</sup> | Also observed in CA3a |
| bFGF | Fibroblast growth factor 2 | Lippoldt et al. <sup>25</sup> |  |
| NT-3 | Neurotrophin-3 | Vigers et al. <sup>26</sup> |  |
| RGS14 | Regulator of G-protein signaling 14 | Lee et al. <sup>27</sup> |  |
| OX<sub>1</sub>R | Orexin receptor 1 | Marcus et al. <sup>28</sup> | Also observed in the crest of the septal DG |
| Cx30.2 | Connexin 30.2 | Kreuzberg et al. <sup>29</sup> |  |
| Nestin | Nestin | Hendrikson et al. <sup>30</sup> |  |
| Ncald | Neurocalcin-δ | Girard et al. <sup>31</sup> |  |
| **Gene expressed in the FC, but not in CA2** |  |  |  |
| Fam163b | Family with sequence similarity 163 member b | Lein et al. <sup>32</sup> | The expression is observed with bare-eyes in the Allen Brain Atlas. Also observed in DG. |
| RgmA | Repulsive guidance molecule A | Schmidtmer and Engelkamp <sup>33</sup> | However, another paper reported the RgmA out of FC <sup>34</sup> |
| **Genes expressed in the CA2, but not in the FC** |  |  |  |
| S100b | S100 protein, beta polypeptide, neural | Dalley et al. <sup>35</sup> |  |

**Table S2.** Primary antibodies used in the current study.

| **Antibodies** | **Host** | **Working dilution** | **Company** | **Catalog number** |
| --- | --- | --- | --- | --- |
| RGS14 | Mouse IgG2a | 1:300 | NeuroMab | #75-170 (RRID: AB_2179931) |
| WFS1 | Rabbit | 1:300 | proteintech | 11558-1-AP |
| NeuN | Mouse IgG1 | 1:500 | millipore | cat.MAB377 |
| NG2 | Rabbit | 1:500 | millipore | cat.AB5320 |
| BrdU | Mouse | 1:300 | Thermo Fisher Scientific | MA3-071 |
| BrdU | Rat | 1:500 | Abcam | Ab6326 |
| Parvalbumin | Rabbit | 1:300 | Swant | PV27 |

**Table S3.** Statistical description of the FC volumetry. The table below show the number of rats in each group and the volume of the FC (unit: mm^3^).

|  | Group | N | Mean | S.E.M. |
| --- | --- | --- | --- | --- |
| T-maze task | Control | 8 | 0.0268 | 0.0019 |
|  | Lesion | 11 | 0.0095 | 0.0011 |
|  | No lesion | 3 | 0.0214 | 0.0034 |
| Spontaneous object exploration | Control | 13 | 0.0267 | 0.0019 |
|  | Lesion | 17 | 0.0125 | 0.0013 |

**Table S4.** Statistical result of the contextual behavior task. Statistically significant results (p<0.05) were denoted with an asterisk. For statistical comparison, repeated measures mixed ANOVA, t-test, paired t-test, Wilcoxon Signed rank test, and Mann-Whitney U test were used and Bonferroni correction is applied to post-hoc test.

| Related figure | Mean ± S.E.M. |  | N | Statistics | Note |
| --- | --- | --- | --- | --- | --- |
| Figure 4d<br>(Lesion vs. control performance) | retrieval, control | 86.3±2.6 | 8 | Surgical condition: F (1, 17) = 1.7, P = 0.207;<br>Learning stage: F (1, 17) = 25.6, P < 0.001*;<br>Surgical condition × Learning stage: F (1, 17) = 5.2, P = 0.036 | lesion vs. control in retrieval<br>P = 0.654 |
|  | retrieval, lesion | 88.9±2.5 | 11 |  |  |
|  | acquisition, control | 75.8±4.0 | 8 |  | lesion vs. control in acquisition<br>P = 0.019* |
|  | acquisition, lesion | 61.3±5.6 | 11 |  |  |
| Ret. D18 | control | 91.6±2.7 | 8 | t (17) = 0.5, P = 0.625 |  |
|  | lesion | 93.2±1.8 | 11 |  |  |
| Acquisition latency | control | 1.93±0.19 | 8 | t (17) = -0.5, P = 0.592 |  |
|  | lesion | 2.05±0.14 | 11 |  |  |
| Related figure | Median (I.Q.R.) |  |  | Statistics |  |
| Figure 4f<br>(trial latency, acquisition) | control, trial 10 | 0.90 (0.42) | 8 | Z = 2.5, P = 0.012 | Wilcoxon Signed rank test |
|  | control, trial 11 | 1.99 (2.73) | 8 |  |  |
|  | lesion, trial 10 | 1.08 (0.67) | 11 | Z = 0.9, P = 0.374 |  |
|  | lesion, trial 11 | 1.28 (0.26) | 11 |  |  |
| Figure 4g (trial latency ratio, trial 10 based) | trial 11, control | 2.08 (0.33) | 8 | Z = 2.4, P = 0.017 | Mann-Whitney U test |
|  | trial 11, lesion | 1.30 (0.85) | 11 |  |  |
|  | trial 12, control | 1.08 (0.36) | 8 | Z = 0.3, P = 0.741 |  |
|  | trial 12, lesion | 0.99 (0.39) | 11 |  |  |

**Table S5.**
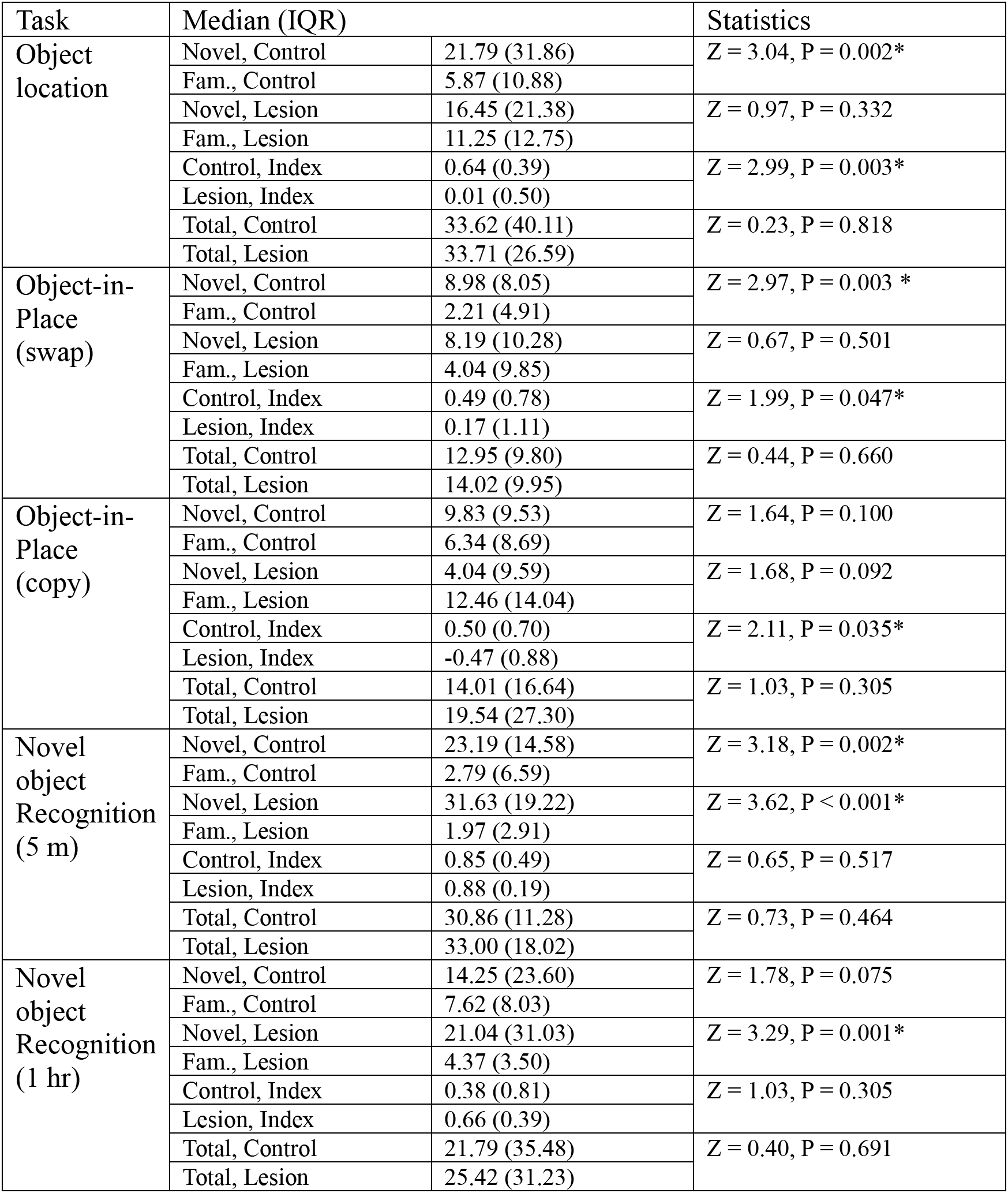
Statistical results of spontaneous object exploration tasks. Asterisks in the p-value column indicate statistical significance. Values represent median (IQR). Wilcoxon signed-rank test was used for the comparison of exploration time, while the Mann-Whitney rank-sum test was used for comparison of the index.

**Table S6.** Statistical result of the electrophysiology experiment (foraging experiment). Asterisks in the p-value column indicate statistical significance. Bonferroni correction was conducted (corrected alpha = 0.166). Values represent Mean ± S.E.M.

| Measurement | FC | dmCA1 | dCA1 | F | df | p-value | FC/dmCA1 | FC/dCA1 | dmCA1/dCA1 |
| --- | --- | --- | --- | --- | --- | --- | --- | --- | --- |
| Spike width (μs) | 312.9±6.8 | 381.9±3.5 | 397.0±5.9 | 40.8 | 2, 311 | P<0.001* | P<0.001* | P<0.001* | P=0.017 |
| Burst index | 21.8±2.2 | 25.0±1.6 | 21.5±2.5 | 1.0 | 2, 311 | P=0.370 | | | |
| Firing rate (sleep, Hz) | 0.47±0.08 | 0.56±0.04 | 0.70±0.07 | 2.8 | 2, 311 | P=0.065 | | | |
| mean firing rate (square field, Hz) | 0.50±0.07 | 0.85±0.06 | 0.89±0.09 | 3.7 | 2, 311 | P=0.025* | P=0.014* | P=0.009* | P=0.643 |
| Peak firing rate (Hz) | 3.86±0.46 | 6.21±0.35 | 6.51±0.481 | 5.4 | 2, 311 | P=0.005* | P=0.003* | P=0.002* | P=0.598 |
| Spatial information (Bits/Spike) | 0.94±0.06 | 1.20±0.05 | 1.23±0.06 | 4.2 | 2, 311 | P=0.016* | P=0.009* | P=0.006* | P=0.632 |
| Spatial coherence | 0.35±0.03 | 0.49±0.02 | 0.53±0.03 | 9.6 | 2, 311 | P<0.001* | P<0.001* | P<0.0001* | P=0.152 |
| Spatial correlation (R, 1<sup>st</sup> half vs. 2<sup>nd</sup> half) | 0.07±0.01 | 0.14±0.01 | 0.16±0.02 | 5.9 | 2,311 | P=0.003* | P=0.005* | P<0.001* | P=0.258 |
| Number of field | 1.45±0.11 | 1.51±0.06 | 1.39±0.07 | 0.9 | 2, 311 | P=0.400 | | | |
| Field size (cm<sup>2</sup>) | 640.9±61.6 | 699.7±33.6 | 759.8±47.6 | 1.1 | 2,455 | P=0.325 | | | |
| In-field firing rate (Hz) | 1.58±0.17 | 2.53±0.12 | 2.78±0.20 | 7.2 | 2,455 | P<0.001* | P=0.001* | P<0.001* | P=0.254 |

**Table S7.** Statistical result of the electrophysiological experiment (SWR). Asterisks in the p-value column indicate statistical significance. Bonferroni correction was conducted (corrected alpha is 0.008 for ripple power comparison and 0.166 for outburst ratio). Values represent Mean ± S.E.M.

| Measurement | SWR power (dB) | SWR outburst ratio |
| --- | --- | --- |
| FC | -35.145±0.499 | 2.234±0.194 |
| dmCA1 | -26.176±0.563 | 11.144±0.747 |
| dCA1 | -26.525±0.346 | 13.805±1.708 |
| CC | -35.922±0.319 | - |
| F | 132.5 | 13.447 |
| df | 3,292 | 2,247 |
| p-value | P<0.001* | P<0.001* |
| Post-hoc test |  |  |
| FC/dmCA1 | P<0.001* | P<0.001* |
| FC/dCA1 | P<0.001* | P<0.001* |
| FC/CC | P=0.384 | - |
| dmCA1/dCA1 | P=0.656 | P=0.088 |
| dmCA1/CC | P<0.001* | - |
| dCA1/CC | P<0.001* | - |

**Table S8.** Statistical result of the electrophysiological experiment (T-maze). Two-way ANOVA is used for statistics. The in-field rate modulation index (RMI) was calculated by each place field (70/55/41 in the dmCA1, 11/14/8 in the FC were analyzed. The number of place fields in the acquisition, retrieval, and shuttling were described, respectively). Values represent Mean ± S.E.M.

| Measurement | Task | dmCA1 | FC | Statistics |
| --- | --- | --- | --- | --- |
| mean firing rate (Hz) | Acq. | 2.53±0.23 | 1.32±0.40 | Subregion: F(1,174)=4.6, P=0.034<br>Learning stage: F(2,174)=1.3, P=0.287<br>Interaction: F(2,174)=0.5, P=0.594 |
| | Ret. | 2.56±0.29 | 1.46±0.35 | |
| | Shuttling | 2.80±0.39 | 2.56±0.88 | |
| Peak Firing rate (Hz) | Acq. | 14.20±1.13 | 7.74±2.66 | Subregion: F(1,174)=6.2, P=0.014<br>Learning stage: F(2,174)=1.3, P=0.285<br>Interaction: F(2,174)=0.9, P=0.393 |
| | Ret. | 13.73±1.25 | 7.80±1.59 | |
| | Shuttling | 14.38±1.51 | 13.63±4.00 | |
| Spatial information (Bits/spk) | Acq. | 1.33±0.08 | 1.13±0.20 | Subregion: F(1,174)=4.1, P=0.04<br>Learning stage: F(2,174)=0.1, P=0.932<br>Interaction: F(2,174)=0.2, P=0.815 |
| | Ret. | 1.25±0.06 | 1.11±0.15 | |
| | Shuttling | 1.36±0.08 | 1.05±0.07 | |
| RMI | Acq. | 0.11±0.01 | 0.16±0.05 | Subregion: F(1,128.4)=4.1, P=0.044<br>Learning stage: F(1,128.4)=0.3, P=0.589<br>Interaction: F(1,128.4)=0.0, P=0.912 |
| | Ret. | 0.12±0.01 | 0.17±0.04 | |
| The normalized firing rate difference | Acq. | 0.37±0.04 | 0.18±0.03 | Subregion: F(1,128)=1.4, P=0.238<br>Learning stage: F(1,128)=1.3, P=0.248<br>Interaction: F(1,128)=1.0, P=0.310 |
| | Ret. | 0.38±0.03 | 0.42±0.08 | |
| | Shuttling | 0.30±0.03 | 0.21±0.04 | |

**Table S9.** Statistical result of the electrophysiological experiment (T-maze). Two-way ANOVA, chi-square, and F-test for variance are used for statistics. Values represent Mean ± S.E.M. For the number of place field maintained until the last trial, Chi-square test is used for a statistical test. The number of episodic fields is 72/62/45 in the dmCA1 and 13/15/10 in the FC. The order is acquisition task, retrieval task, and shuttling session, respectively. Despite significant interaction in the analysis of place field duration, we did not find any difference between tasks in the FC.

| Measurement | Task | dmCA1 | FC | Statistics |
| --- | --- | --- | --- | --- |
| Episodic field start trial | Acq. | 16.63±3.56 | 27.15±9.10 | Subregion:<br>F(1,178.2)=6.5,<br>P=0.011<br>Learning stage:<br>F(2,176.9)=1.9,<br>P=0.153<br>Interaction:<br>F(2,176.9)=1.7,<br>P=0.181 |
|  | Ret. | 31.40±5.44 | 37.33±11.58 |  |
|  | Shuttling | 17.78±4.60 | 52.50±15.44 |  |
| The number of place (ceased/ maintained/ total) | Acq. | 13/59/72 | 8/5/13 | Subregion: $\chi^2(1)$=12.3,<br>P<0.001<br>Learning stage:<br>$\chi^2(2)$=1.8, P=0.408<br>Interaction: $\chi^2(2)$=1.8,<br>P=0.413 |
|  | Ret. | 14/48/62 | 6/9/15 |  |
|  | Shuttling | 14/31/45 | 6/4/10 |  |
| Place field duration | Acq. | 55.56±2.25 | 28.43±3.28 | Subregion:<br>F(1,180)=23.0, P<0.001<br>Learning stage:<br>F(2,180.3)=0.3,<br>P=0.712<br>Interaction:<br>F(2,180.3)=3.1,<br>P=0.048† |
|  | Ret. | 45.08±2.61 | 39.67±5.65 |  |
|  | Shuttling | 48.25±2.80 | 29.83±4.49 |  |
| †Post-hoc test: FC, Ret. vs. FC, Acq.: P=0.132; FC, shuttling vs. FC, Acq.: P=0.821; FC, shuttling vs. FC, Ret: P=0.241 |  |  |  |  |

